# Visual mental imagery in typical imagers and in aphantasia: A millimeter-scale 7-T fMRI study

**DOI:** 10.1101/2023.06.14.544909

**Authors:** Jianghao Liu, Minye Zhan, Dounia Hajhajate, Alfredo Spagna, Stanislas Dehaene, Laurent Cohen, Paolo Bartolomeo

## Abstract

Most of us effortlessly describe visual objects, whether seen or remembered. Yet, around 4% of people report congenital aphantasia: they struggle to visualize objects despite being able to describe their visual appearance. What neural mechanisms create this disparity between subjective experience and objective performance? Aphantasia can provide novel insights into conscious processing and awareness. We used ultra-high field 7T fMRI to establish the neural circuits involved in visual mental imagery and perception, and to elucidate the neural mechanisms associated with the processing of internally generated visual information in the absence of imagery experience in congenital aphantasia. Ten typical imagers and 10 aphantasic individuals performed imagery and perceptual tasks in five domains: object shape, object color, written words, faces, and spatial relationships. In typical imagers, imagery tasks activated left-hemisphere frontoparietal areas, the relevant domain-preferring areas in the ventral temporal cortex partly overlapping with the perceptual domain-preferring areas, and a domain-general area in the left fusiform gyrus (the Fusiform Imagery Node). The results were valid for each individual participant. In aphantasic individuals, imagery activated similar visual areas, but there was reduced functional connectivity between the Fusiform Imagery Node and frontoparietal areas. Our results unveil the domain-general and domain-specific circuits of visual mental imagery, their functional disorganization in aphantasia, and support the general hypothesis that conscious visual experience - whether perceived or imagined - depends on the integrated activity of high-level visual cortex and frontoparietal networks.

## Introduction

Visual mental imagery is the ability to experience visual information in the absence of actual external stimuli. However, approximately 4% of individuals report experiencing weak or absent visual mental imagery (Dance et al., 2022), a condition known as congenital aphantasia (Zeman et al., 2015). The nature of aphantasia remains elusive. Some perspectives suggest that aphantasia is a deficit in introspection or awareness of internal visual imagery (Liu & Bartolomeo, 2023), while others propose it as a deficiency in voluntarily-generated mental imagery (Zeman et al., 2015), or in general mental imagery capacities (Keogh & Pearson, 2018). Surprisingly, individuals with congenital aphantasia can typically provide accurate answers from memory to questions on the visual appearance of objects, despite the absence of subjective imagery experience. For example, these individuals can correctly indicate which fruit is darker red between strawberries or cherries (Liu & Bartolomeo, 2023), and perform many imagery-related tasks such as visual working memory and mental rotation (Pounder et al., 2022). Thus, aphantasia provides a unique window into the mechanisms underlying conscious processing and awareness. Why don’t aphantasic individuals experience visual mental images?

Models of visual mental imagery are structured around two primary components: the frontoparietal (FP) cortex and the visual cortex. However, models diverge on the specific portions of the visual cortex crucial for visual mental imagery. While some emphasize the importance of early visual areas (EVA)(Pearson, 2019), others highlight the role of the high- level visual areas in the ventral temporal cortex (VTC)(Spagna et al., 2021). These competing models make different predictions on the possible neural underpinnings of aphantasia.

Many neuroimaging studies predominantly focused on the primary visual cortex (Kosslyn et al., 2006). For example, individual vividness was related to the activation level of the V1 (Cui et al., 2007) and imagined stimuli could be decoded or reconstructed from the representational pattern of the EVA (Naselaris et al., 2015; Senden et al., 2019; Thirion et al., 2006), possibly driven by the signal from the deep cortical layers of V1 (Bergmann et al., 2024). On the other hand, evidence from acquired brain lesions in neurological patients demonstrating preserved imagery abilities after lesions to the EVA or their connections (Bartolomeo et al., 1998, 2020) strongly suggests that the EVA are not necessary for visual mental imagery. If, however, visual mental imagery experience does rely on EVA activity, then this activity should be dysfunctional in aphantasia.

A recent meta-analysis of 27 imagery fMRI studies (Spagna et al., 2021) highlighted imagery-related activity in FP regions and in a specific region of the left hemisphere fusiform gyrus, which was active independent of the imagery domain. Spagna et al. labeled this region the Fusiform Imagery Node (FIN). In line with the localization of the FIN, lesion studies revealed domain-general imagery deficits following extensive damage to the left temporal lobe (Bartolomeo et al., 2002; Liu et al., 2022; Moro et al., 2008; Thorudottir et al., 2020). Nevertheless, no study has yet examined the functional properties of the putative FIN during visual mental imagery, or its possible dysfunction in aphantasia.

A crucial yet understudied aspect is the role of domain-preferring VTC cortical patches in visual mental imagery. The VTC contains cortical patches with relatively selective activity for specific perceptual domains, such as faces, words, and colors (Cohen et al., 2000; Kanwisher et al., 1997; Lafer-Sousa et al., 2016). Early fMRI studies showed that imagery of faces and places reactivated the fusiform face area (FFA) and the parahippocampal place area (PPA), respectively (Ishai et al., 2000; O’Craven & Kanwisher, 2000). Also, color information can be decoded from a color-biased region during color imagery (Bannert & Bartels, 2018). Lesion studies demonstrated that brain damage to the temporal cortex (but not to the occipital cortex) can result in deficits in imagery that are specific to particular visual domains, such as object shape, object color, written words, faces, and spatial relationships (Bartolomeo, 2002; Goldenberg, 1993). These findings underscore the necessity for a more systematic investigation of domain-preferring cortex in visual mental imagery. Do these regions contribute to imagery vividness in typical imagery or to its absence in aphantasia?

Finally, a further open question concerns the role of FP networks. Clinical (Bartolomeo, 2007), neuroimaging (Chica et al., 2012) and neurophysiological (Liu et al., 2023; Spagna et al., 2022) evidence highlights the crucial role of FP network activity in conscious perception. Can FP dysfunction result in the lack of experiential correlates of access to offline visual information in aphantasia? This could be characterized by an alteration of functional brain networks akin to those observed in some neurodevelopmental disorders (Sokolowski & Levine, 2023). For example, Milton et al (2021) reported reduced resting-state functional connectivity between prefrontal areas and the visual cortex in aphantasic participants, compared to individuals reporting high imagery vividness. However, it remains uncertain whether aphantasia displays impaired connectivity during attempted mental imagery.

Neuroimaging studies have encountered challenges in distinguishing between domain-general and domain-specific mechanisms in visual mental imagery for the following reasons: i) the difficulty in demonstrating a causal link between activations (e.g. in EVA) and imagery processes (Bartolomeo et al., 2020), ii) the dearth of studies that examine several domains with naturalistic stimuli, which are essential for eliciting activity in high-level visual cortex including both the FIN and domain-preferring areas, and iii) the limited spatial resolution of conventional 3T fMRI, along with techniques like group averaging, which hindered the ability to distinguish closely packed domain-preferring visual regions (Saxe et al., 2006). For instance, the face, word, and body-preferring areas in the VTC are situated in close proximity (Grill-Spector & Weiner, 2014). Moreover, given the individual variability in the topographical organization of the VTC (Conway, 2018; Grill-Spector & Weiner, 2014), assessing the degree of overlap of activations between imagery and perception requires the use of methods with high spatial resolution in individual participants. Compared to functional neuroimaging, lesion studies possess causal power (Bartolomeo et al., 2020), but are affected by limitations including limited spatial resolution, disruption of white-matter connections beyond the gray-matter lesions, and lesion-induced plasticity or reorganization.

Here, we aim to i) identify the neural circuits involved in domain-general and domain- specific voluntary imagery in individual participants; ii) compare them with those engaged in visual perception; iii) examine the neural processes associated with accurate information processing in the absence of experiential correlates in aphantasia. We circumvent the above-mentioned limitations by i) using ultra-high field 7-Tesla fMRI during tightly matched imagery and perceptual tasks in five visual domains suggested by lesion studies (Goldenberg, 1993): object shape, object color, written words, faces, and spatial relationships, ii) studying typical imagers and individuals with aphantasia. Crucially, our tasks involve visualizing real-world stimuli retrieved from long-term memory, without any visual cues. We predicted that, under the EVA hypothesis, typical imagers would exhibit normal activities including EVA activation, representational content and connectivity, whereas those with aphantasia would demonstrate dysfunctional EVA activity or connectivity. Alternatively, aphantasia could be associated with abnormal activity or connectivity of high-level visual cortex, either in the domain-general or domain-preferring VTC, as suggested by lesion localization in neurological patients with impaired visual mental imagery.

## Method

### Participants

Ten typical imagers (mean age ± SD, 29.28 ± 8.47, 6 female) were recruited from the CNRS RISC volunteer database (https://www.risc.cnrs.fr/). All typical imagers had average or high vivid mental imagery with Vividness of Visual Imagery Questionnaire (VVIQ) score greater than 55 (mean ± SD, 71.70 ± 7.07, out of a total score of 80). Ten individuals with congenital aphantasia (mean age ± SD, 28.69 ± 8.27, 6 female) were recruited from French-language groups on aphantasia in various online social media. All aphantasic participants reported a *complete* life-long inability to generate visual mental imagery, and this was confirmed by an individual interview with author D.H. (a clinical psychologist) during the recruitment. All aphantasic individuals reported scoring 16 out of 80 (“no image at all” for all questions) to the VVIQ. Interestingly, many aphantasics (8 out of 10) confirmed that they tend to perceive pictures with a *divide-and-label* strategy, akin to semantic conversion from visual features into content lists. While asked to visualize, they briefly recall the list of semantic labels to respond to the relevant question. Notably, they reported difficulties in remembering complex or unfamiliar pictures, which might reflect lower efficiency of the semantic encoding strategy in this case. All participants were right-handed with normal or corrected-to-normal vision and had no history of neurological/psychiatric disorders. Aphantasic participants did not differ from typical imagers in age (t = 0.35, p = 0.73, Cohen’s d = 0.16) or in education level (t =1.33, *p* = 0.20, Cohen’s d = 0.62 ). Participants provided written consent before the study and received monetary compensation after the study. The study was approved by CEA and, according to French bioethical law, by a randomly selected regional ethical committee for biomedical research (CPP 100055 to NeuroSpin center) and the study was carried out in accordance with the declaration of Helsinki.

### Introspective reports

Before the fMRI session, participants completed the French versions of the VVIQ (Santarpia et al., 2008) and Object-Spatial Imagery Questionnaire (OSIQ) (Blajenkova et al., 2006) questionnaires to assess the subjective vividness of their visual mental imagery. The OSIQ consists of two scales assessing preferences for representing and processing of object imagery about pictorial and high-resolution images of individual objects, and of spatial imagery about semantic images and spatial relations amongst objects. The results confirmed the drastic reduction of imagery ability in aphantasic individuals compared to typical imagers (VVIQ, BF = infinity, Cohen’s d = 9.66; OSIQ object, BF = 2.06×10^9^, Cohen’s d = 7.54), but no reduction in spatial ability (OSIQ spatial: BF = 0.46, Cohen’s d = 0.29).

### Stimuli and fMRI experimental design

In the scanner, participants performed a longer version of the enhanced Battérie Imagerie- Perception - eBIP (Liu & Bartolomeo, 2023). The current version of the battery assesses (1) imagery of object shapes (Fig. 1A), object colors (Fig. 1B), faces (Fig. 1C), letters (Fig. 1D) and spatial relationships on an imaginary map of France (Fig. 1E); (2) a non-imagery control task using abstract words (Fig. 1F), and (3) an audio-visual perception task using the same items as in the imagery tasks (Fig. 1G). In the imagery tasks, participants heard a word indicating a particular imagery domain (e.g., “shape”), followed by 2 words, designating the items the participant is required to imagine (e.g. “beaver”, “fox”). They were instructed to generate and maintain mental images as vivid as possible for each item. Eight seconds after the second items, they heard an attribute word (e.g. “long”). They then pressed one of two buttons indicating which of the items is most closely associated with the attribute (e.g. which of the animals they associate with the attribute “long”, see Fig. 1A). Finally, they reported the overall vividness of their mental imagery in that trial on a 4-level Likert scale by pressing one of 4 buttons of an MR-compatible button box (Current Designs, Philadelphia, USA), where button 1 indicated “no image at all” and button 4 indicated a “vivid and realistic image”. In the shape imagery task, participants had to decide which item was longer or rounder. In the color imagery task, participants had to decide which fruit or vegetable had darker or lighter color. In the letter imagery task, participants had to imagine the shape of French words in lowercase and had to decide which word had ascenders (e.g., t, l, d) or descenders (e.g. j, p, y). In the famous-faces imagery task, participants had to decide which celebrity had a more round or oval face. In the map-of-France imagery task, participants had to decide which city was located to the left or the right to Paris. In the non-imagery abstract-word task, participants had to decide which of two abstract words (e.g., “routine”, convention”) was semantically closer to a third word (e.g. “society”). While visual imagery cannot be completely excluded in this setting (or in any other settings), we selected abstract words for this task in order to minimize its engagement. In the perception task, the same stimuli used for the imagery tasks were presented in an audio-visual format. In the abstract-word and the perception tasks participants rated their confidence on a 4-level Likert scale, instead of rating vividness. The auditory stimulus were voice recordings of the corresponding French names per item. All voice recordings were generated in an online TTS engine (https://texttospeechrobot.com/; fr-FR_ReneeVoice) and digitized at a 44.1 kHz sampling rate. The images shown in the perception task were color photographs of the corresponding items on a gray background. The words in letters perception task were rendered in “Monotype Corsiva” font to display words because in pilot testing some participants reported that this font was more natural and closer to their everyday visual experience. The resulting occipito-temporal activations in fMRI (-49, -59, -7; left fusiform; 1,989 mm3) were in line with the extensive neuroimaging literature on reading.

**Fig 1.**
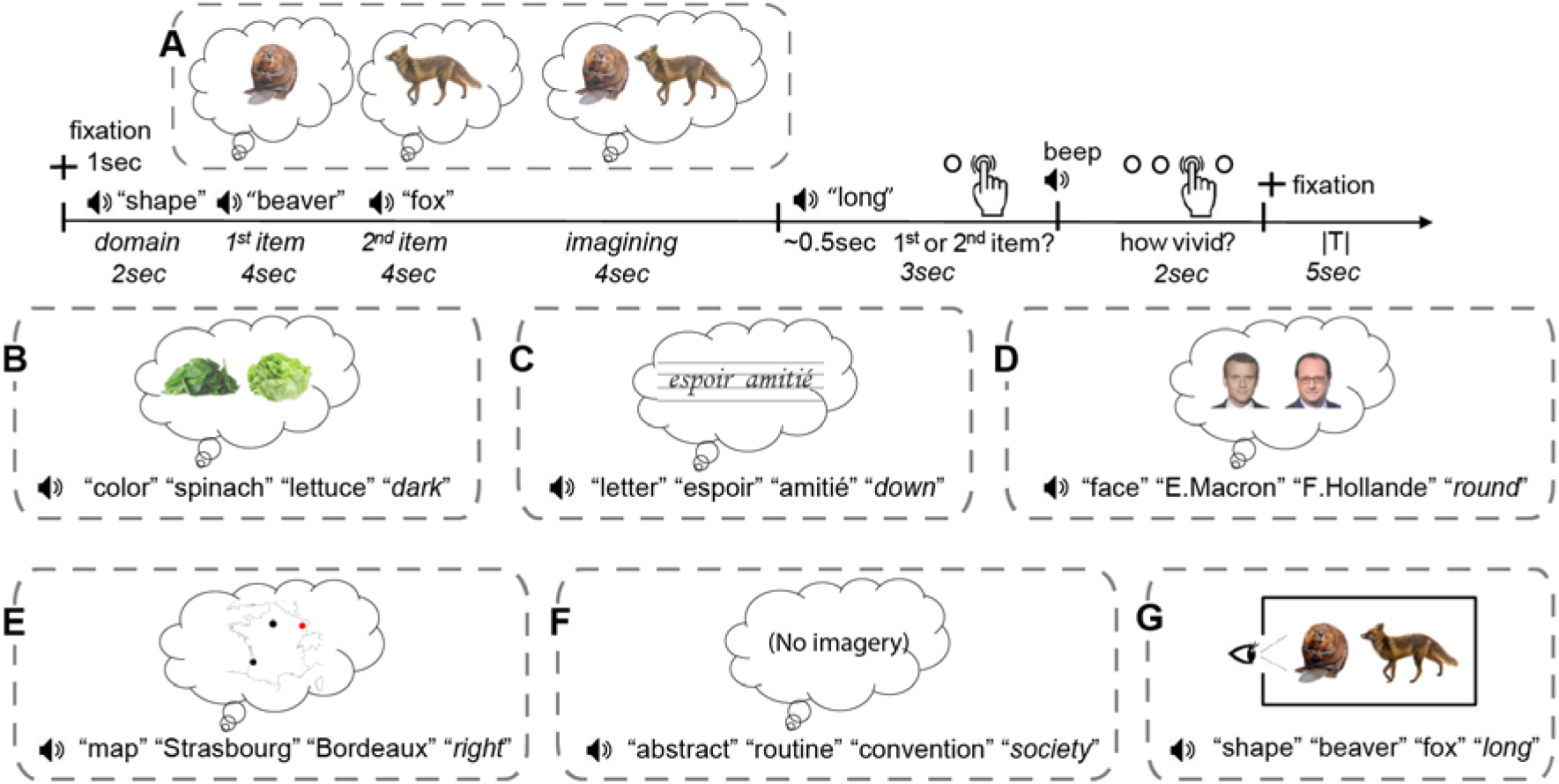
Imagery and Perceptual Tasks. Examples of trials of the eBIP. The eBIP comprises imagery tasks of five domains: shapes, colors, letters, faces, and spatial relationships (A-E), a control task with abstract words (F), and a perception task (G) in five audio-visual domains using the same items as in the imagery tasks. In the imagery tasks, participants heard a word indicating a particular imagery domain (e.g., “shape”), followed by 2 words (e.g. “beaver”, “fox”). Participants were instructed to generate and maintain mental images as vivid as possible for each of these 2 words. Eight seconds after the second item, participants heard an attribute word (e.g. “long”). They then pressed one of two buttons indicating which of the items was most closely associated with the attribute (e.g. which of the animals they associate with the attribute “long”). In the shape imagery task, participants had to decide which of the 2 words designated a longer or rounder item. In the color imagery task, participants had to decide which fruit or vegetable had darker or lighter color. In the letter imagery task, participants had to imagine the shape of French words in lowercase and had to decide which word had ascenders (e.g., t, l, d) or descenders (e.g. j, p, y). In the famous faces imagery task, participants had to decide which of 2 named celebrities had a more round or oval face. In the map-of-France imagery task, participants had to decide which of 2 cities was located to the left or the right of Paris. In the non-imagery abstract word task, participants had to decide which of two abstract words (e.g., “routine”, convention”) was semantically closer to a third word (e.g. “society”). In the perception task, the same stimuli used for the imagery tasks were presented in an audio-visual format.

### Data acquisition

The brain images are acquired using an ultra-high field 7-Tesla Magnetom scanner (Siemens, Erlangen, Germany) with a 1Tx/32Rx head coil (Nova medical, Wilmington, USA) at the NeuroSpin center of the French Alternative Energies and Atomic Energy Commission (CEA). No dielectric pads were used due to the limited space within the head coil with the headphone (OptoActive II, Optoacoustics, Israel). To minimize light reflections inside the head coil, a piece of black paper was inserted to cover the inner surface of the transmitter coil element. Stimuli were presented via a BOLDscreen 32 LCD screen (Cambridge Research Systems, UK, 69.84 x 39.29 cm, resolution=1920 x 1080 pixels, refresh rate=120 Hz, viewing distance ∼200 cm), at the head-end of the scanner bore. Participants viewed the screen through a mirror attached to the head coil.

The fMRI scan session lasted 150 minutes in total, split into 5 runs. The first 3 runs consisted of 90 imagery-only trials (30 trials per run, each including 6 trials per domain, 12 min 14 s per run, 367 volumes with 2 volumes of resting at the beginning of each run). The absence of visual perceptual trials prevented potential spillovers of perceptual responses to imagery trials. The last 2 runs consisted of 36 trials each (6 abstract word trials, 15 imagery trials and 15 audio-visual perception trials per run, each including 3 trials per domain, 14 min 38 s per run, 439 volumes), in that order. Notably, items in the imagery trials and perceptual trials of the last two runs are identical. The inter-trial intervals were jittered between 3s and 7s. Importantly, the order of trials within each task type was fully randomized across domains. For each imagery domain, the order of items was counterbalanced, i.e., each item is presented once as the first item and once as the second item. Participants were instructed to keep their eyes open during scans and to pay attention to the fixation on a gray screen displayed at the beginning of each trial and during the intertrial period. During the imagery and abstract word tasks, they saw an empty gray screen without fixation cross. During the perception tasks, they saw visual stimuli while simultaneously hearing the corresponding auditory stimuli. The perception tasks also enabled the localization of domain-preferring regions.

Functional data were acquired with a 2D gradient-echo EPI sequence (TR = 2000 ms, TE = 28 ms, voxel size = 1.2 mm isotropic, multiband acceleration factor=2; encoding direction: anterior to posterior, iPAT=3, flip angle = 75, partial Fourier=6/8, bandwidth=1488 Hz/Px, echo spacing=0.78 ms, number of slices=70, no gap, reference scan mode: GRE, MB LeakBlock kernel: off, fat suppression enabled). The slab is tilted upwards to cover most parts of the brain except the anterior temporal lobe, and to avoid the eyeballs in the slab. To correct for EPI distortion, a 5-volume functional run with the opposite phase encoding direction (posterior to anterior) was acquired immediately before each task run. Participants were instructed not to move between these pairs of two runs. Manual interactive shimming of the B0 field was performed for all participants. The system voltage was set to 250 V for all sessions, and the fat suppression was decreased per run to ensure the specific absorption rate for all functional runs did not surpass 62%. To minimize artifacts and increase signal-to- noise ratio around the ventral temporal cortex, the functional data acquisition slab was tilted in a way that excluded the eyes and the ear canal signal dropout region, so that the ventral temporal cortex especially the anterior occipital-temporal sulcus above the ear canal was covered. However, part of the anterior temporal lobe was not able to be included (Fig.S2A).

High-resolution MP2RAGE anatomical images were obtained between the third and the fourth functional runs (resolution=0.65 mm isotropic, TR=5000 ms, TE=2.51 ms, TI1/TI2=900/2750 ms, flip angles=5/3, iPAT=2, bandwidth=250 Hz/Px, echo spacing=7 ms).

### Behavioral data analysis

Our behavioral data analysis was similar to that of a study using the same tasks with a more extensive cohort of 117 participants (Liu & Bartolomeo, 2023). We conducted two Bayesian repeated measures ANOVAs within each modality (imagery, perception), with the factors of Group (Aphantasia, Typical imagers) and Domain (Shape, Color, Word, Face, Space). The dependent variables for the imagery tasks were accuracy (arcsine-transformed proportions of correct responses); response times (RTs) and trial-by-trial vividness scores (translated to a 0-1 scale and arcsine-transformed), and for the perceptual tasks were accuracy, RTs and trial-by-trial confidence scores (translated to a 0-1 scale and arcsine-transformed). For each participant, we excluded trials with response times (RT) faster than 150 ms or exceeding three SDs from the participant’s mean. Statistical tests were performed using JASP 0.16.2 (https://jasp-stats.org/), and used the JASP default priors. A commonly accepted convention is that Bayes factors (BF10 or BFs) between 3 and 10 indicate moderate evidence in favor of the model in the numerator (H1); BFs between 10 and 30 indicate strong evidence; BFs larger than 30 indicate very strong evidence. The inverse of these cut-offs values provides moderate (0.33-0.1), strong (0.1-0.03), or very strong evidence (<0.03) for the model in the denominator (H0), i.e. the null hypothesis. We adopted the default setting in JASP for the a-priori values, which uses a uniform distribution for candidate models, e.g., P(M) = 0.5 for two alternative models.

### fMRI data Preprocessing

We processed the fMRI data with BrainVoyager (Version 22.0.2.4572, Brain Innovation, Maastricht, The Netherlands, https://www.brainvoyager.com/ ), MATLAB (version R2018b), and NeuroElf 1.1 toolbox (http://neuroelf.net/) implemented in MATLAB. The functional data underwent EPI distortion correction using the posterior-to-anterior directional functional volumes (COPE plugin in BrainVoyager), where the in-plane voxel displacement map between the actual task run and the first volume of the preceding distortion correction run (in reversed phase encoding direction) was computed, and applied to the task run. The distortion-corrected data was then corrected for slice scan time (sinc interpolation, slice scanning order read from the slice time table in the DICOM headers), 3D rigid motion correction (trilinear for estimation, sinc for applying the correction, aligned to the first volume within each run), high-pass temporal filtering (GLM with Fourier basis set, number of cycles=3). No spatial smoothing was applied to the data at this stage.

The MP2RAGE anatomical data consisted of four image types: inversion 1, inversion 2, quantitative T1, uniform. To have a similar appearance to the conventional MPRAGE anatomical data, the uniform image was divided by the T1 image (an optional step), and the background noise was masked out by the inversion 2 image. The resulting anatomical image was resampled to 0.6 mm isotropic (framing cube dimension: 384×384×384), and transformed into Talairach space (TAL). All the coordinates reported in our manuscript are Talairach coordinates. For data visualization, the white matter-gray matter boundary was segmented in TAL space, and reconstructed as surface meshes.

For fMRI across-run co-registration, the fourth functional run was co-registered to the anatomical data, then all the other functional runs were manually co-registered to the fourth functional run. The across-run co-registration quality was inspected visually with animations looping through the first volumes across runs in TAL space. In traditional fMRI-to-anatomy coregistration, the contrast and resolution of the fMRI (T2*) is very different from the anatomical (T1) images (see Fig. S2A), and often includes imperfections such as EPI distortion (potentially not fully corrected by top-up distortion correction). In that scheme, the fMRI-to-anatomy coregistration algorithm is not perfect (especially with partial brain coverages, or the acquisition slab moved too much e.g. across different sessions), and would induce machine errors everytime the coregistration is performed. In comparison, our fMRI-to-fMRI coregistration approach minimizes the difference of image contrasts between runs (always T2* and always the same resolution), and allows both machine and human coregistration and quality assurance with high precision. For the choice of the 4th fMRI run, participants initially performed the first three runs, then the T1 anatomical scan, and then the 4th and 5th runs. The first TR functional image of the 4th run was closest to the anatomical scan, where participant’s inter-run head movements (e.g. rest and stretching between runs, plus the within-run head movements accumulated across the whole scanning session) would be the smallest. After the quality checks, all functional images were then transformed into TAL space.

After alignment across runs, the functional data of the main experiment were then spatially smoothed with 6 mm FWHM for group-level univariate analysis due to the individual anatomical/functional variability. No spatial smoothing was applied in the case of individual analysis. All results in this study were computed in the volume space.

### Univariate analysis

Two different general linear models (GLMs) were defined with the following main predictors: 1) five domains in imagery-only trials (5 predictors in total) in the first 3 runs; 2) abstract word and five domains in imagery and in perception trails (11 predictors in total) in the last 2 runs.

The period of predictors started from the first domain words to the attribute word before rating periods. For all the GLM models above, the time courses were %-transformed, the main predictors were convolved with a two-gamma hemodynamic response function, and the 6 parameters of participant’s head motion were z-scored and entered as confounding factors. AR(2) correction was used for correcting serial correlations. The two GLM models were applied to both no-smoothed fMRI datasets for individual analysis and 6 mm FWHM smoothed fMRI datasets for group-level analysis.

The contrast of single imagery domain *versus* other four domains was conducted with trials in run 1-3 to identify domain-preferring regions for each domain. The contrast of all Imagery *versus* Abstract words and single perceptual domain *versus* other four domains was conducted with trials in run 4-5.

The group random-effect GLM analysis was performed for each predictor set with smoothed data. Cluster size thresholds for all group-level contrast maps in this study were estimated using Monte-Carlo simulation (alpha level=0.05, numbers of iterations=5000, initial p<0.005), with the BrainVoyager plugin Cluster-Level Statistical Threshold Estimator (https://support.brainvoyager.com/brainvoyager/functional-analysis-statistics/46-tresholding-multiple-comparisons-problem/226-plugin-help-cluster-thresholding), masked with the common functional data coverage across 10 participants in each group. We used a p-value of 0.005 instead of 0.001 as a necessary compromise between individual variability i.e. false negative results, and the risk of false positive results. Our 7T fMRI data at 1.2 mm isotropic resolution is highly robust at the individual-subject level because of the high image contrast between white/gray matters and temporal signal-to-noise ratio (tSNR), but it has also exacerbated inter-individual anatomical and functional differences. Besides, the Monte-Carlo clustering method does not overly inflate the activated cluster sizes under different initial p values in our data. For example, we made 3 Monte Carlo simulations under initial p=0.01, and 0.001 for the contrast of all Imagery versus Abstract words with 5,000 iterations, and observed consistent surviving clusters across different thresholds.

### Delineating individual functional foveal V1 and pheripheral V1

Due to the substantial functional variability across participants in EVA (Benson et al., 2022),, we mapped the location of the foveal V1 and the pheripheral V1 in each individual based on brain anatomy and functional activation during perception. For peripheral V1, the individual brain was first co-registered to a functional visual atlas visfAtlas (Rosenke et al., 2021) with linear transformation to each individual’s brain anatomy through meticulous manual alignment and thorough visual inspection, particularly in the vicinity of the calcarine sulcus. The foveal V1 was manually delimited in the individual brain in the lateral occipital lobe around the retro calcarine sulcus. The activity profiles of both V1 subregions confirmed its activation during perception.

### Delineating EVA, ventral and dorsal visual pathway ROIs

To account for the variation in the location of the activation in different domains, we visually delineated the EVA, ventral and dorsal visual pathway ROIs. The EVA consists of cuneus, the posterior part of lingual gyrus/pericalcarine. The ventral visual pathway ROI consists of anterior part of lingual gyrus, fusiform gyrus and inferior temporal gyrus. The dorsal visual pathway ROI consists of superior occipital cortex, inferior parietal lobule and posterior part of superior parietal lobule. We then counted the voxels in the volumetric intersection of domain- specific activation in the EVA, ventral, and dorsal ROIs, respectively, in each domain.

### Representational similarity analysis (RSA)

To assess the group difference in representational information during imagery, we performed RSA analysis within all imagery domain-general regions activated in the contrast of all Imagery *versus* Abstract words, and additionally within V1. We obtained voxel-wise % signal change from TR 4-5 (normalized by the TRs -2 to 0 as the baseline), corresponding to the maximum activation of the first item and with minimal influence from the second item in each trial. For imagery similarity analysis, we obtained brain activity from 90 imagery trials (18 per domain) in run 1-3. For perceptual similarity analysis, we extracted brain activity from 30 perceptual trials (6 per domain) in run 4-5. In imagery-perceptual similarity comparison, we used brain activity from imagery trials in run 4-5, which had the same items as in perceptual trials. We calculated the correlation between the multivoxel patterns of each pair of stimuli using the Pearson correlation coefficient to generate a representational dissimilarity matrix (RDM). Before comparing between groups, we first assessed within-group variability by calculating the correlation between each participant’s RDM and the mean RDM derived from the remaining individuals within the same group. For either group, there was no evidence of any differences in within-group consistency of these regions (all BFs < 2.51).

### Psychophysiological interactions (PPIs) analysis

To investigate whole-brain task-specific functional connectivity, we conducted PPI analysis with trials in run 4-5. We built each PPI design matrix by 1) generating a task contrast regressor of two conditions, balanced by a regressor of the sum of two conditions, both convolved with the hemodynamic response function (HRF); 2) extracting a demeaned time course from the seed ROI; 3) generating an interaction regressor as an element-by-element product of the HRF-convolved task contrast and seed ROI regressors; 4) adding the 6 parameters of participant’s z-scored head motion as confounding factors. Specifically, to investigate domain-general connectivity, we grouped all five imagery domains as a single main condition and all five perception domains as another main condition, and included a separate main condition for Abstract words. In each PPI contrast, a task contrast regressor was established, such as the contrast between all Imagery *versus* Abstract words, which was defined by subtracting the main condition for Abstract words from the main condition for Imagery. On domain-specific connectivity, we built the task contrast regressor by subtracting the Abstract words condition from single domain condition, while keeping the other imagery/perceptual domain conditions unchanged. In BrainVoyager, we conducted GLMs using the PPI design matrix for each contrast separately on fMRI datasets that were smoothed with 6 mm FWHM. The resulting group-level maps were thresholded at p<0.005 for cluster size correction.

### Task-residual functional connectivity analysis

In order to identify putative direct upstream or downstream areas within the same functional structure of the FIN in the task, we further performed functional connectivity using task residual data that exploits the variance remaining after removing the mean task-related signal from a time series. We smoothed the task data of the run 1-3 at 6 mm FWHM, regressed out the task-related activity by deconvolution analysis (12 stick predictors per stimulus, covering the evolvement of the BOLD shape per trial) and head motion parameters. Using individual FIN as the seed region, we averaged the extracted residual time course across voxels, and correlated it with the residual time courses of all voxels in each run, resulting in one correlation R map per run. The connectivity pattern across the runs was stable for all participants. The R maps were Fisher’s Z-transformed, averaged across runs per participant for group-level comparisons. The resulting group-level maps were thresholded at p<0.005 for cluster size correction using Monte-Carlo simulation (alpha level=0.05, numbers of iterations=5000, initial p<0.005).

### Individual trial-by-trial parametric modulation of vividness

We fitted individual GLM with trial-by-trial vividness ratings as an main predictor, applied to data from the run 1 to 3 with 6 mm FWHM smoothed fMRI datasets. There were 90 imagery trials in total to estimate the individual parametric modulation of vividness.

## Results

We first present the behavioral results, then the fMRI findings in the domain-preferring VTC areas, in the domain-general FIN, and finally in the EVA. For each region, we compare perception vs. imagery, and typical imagers vs. aphantasic participants, using univariate and multivariate methods, plus the study of functional connectivity. The high spatial resolution of 7T fMRI allowed us to clearly observe domain-specific BOLD responses in the ventral temporal cortex of individual subjects. As a consequence, we adopted a multiple single case approach, and we included results of individual participants whenever possible.

### Behavioral results

For typical imagers, the average trial-by-trial vividness score was 3.52 on a scale of 1 to 4, while for aphantasic individuals the average score was 1.11 (BF = 5.387×10^11^, see Fig. S1B for all behavioral results). Nevertheless, aphantasic individuals exhibited comparable levels of accuracy to typical imagers in both imagery (Bayesian repeated measures ANOVA with the factors of Group x Domain, the main Group effect BF = 0.25 without interaction, partial eta-squared = 0.008) and perception (BF = 0.26 without interaction, partial eta-squared = 0.002). Their RTs were 0.21s slower on the imagery tasks and 0.43s slower on the perception tasks (BF = 30 and 85, respectively, partial eta-squareds > 0.28), and they had lower confidence in their responses on perceptual tasks (BF = 3.59, partial eta-squared = 0.17), consistent with similar findings from a more extensive cohort of 117 participants (Liu & Bartolomeo, 2023). Importantly, however, there was no evidence of group difference on either accuracy (BF = 0.46), RTs (BF = 0.41) or confidence scores (BF = 1.63) in the abstract words task, suggesting that the group difference was specific to visual items.

### Domain-preferring activations overlap during imagery and perception

In both imagery and perception, we localized domain-preferring regions by contrasting activity elicited by each domain minus the other four domains in order to answer three questions: Does local domain-preference prevail in imagery as it does in perception? Are domain-preferring regions the same in imagery and in perception? Do typical and aphantasic participants differ in those respects? There was substantial individual variability in the location of activations, and each domain could activate multiple cortical patches, including in the VTC, consistent with previous 3T and 7T studies with single-subject analyses (Zhan et al., 2023; Zhen et al., 2015). For example, face-preferring fusiform patches showed substantial individual variability during perception (Fig. S3B). Thus, we report individual domain-preferring maps with a summary of regions for each group (each domain *versus* the other four domains, thresholded at p<0.001 uncorrected for all individual maps, no data smoothing, and cluster size > 12 voxels).

**<u>In typical imagers</u>**, during **imagery tasks** (Fig. 2A displays the results for a representative typical imager and see Fig. S3 for other individual activation maps; see Table 1 for a summary), i) shape imagery activated the fusiform gyrus (FG, 5 left and 2 bilateral out of 10 participants) and lateral occipital complex (LOC, 3 right, 2 bilateral); ii) color imagery activated medial FG (6 left, 1 right, 2 bilateral), parahippocampal gyrus (PHG, 4 left, 1 right, 3 bilateral), posterior occipitotemporal sulcus (OTS, 4 left, 6 bilateral), orbitofrontal cortex (OFC, 2 left, 7 bilateral), inferior frontal sulcus (IFS, 2 left, 7 bilateral); iii) word imagery activated posterior OTS (2 left, 1 right, 7 bilateral), lateral occipitotemporal cortex (LOTC, 4 left, 4 bilateral), dorsolateral prefrontal cortex (dlPFC) and Intraparietal sulcus (IPS); iv) face imagery activated OTS (FFA, 2 left, 7 bilateral), middle superior temporal gyrus (mSTG, 2 left, 6 bilateral), ventral posterior cingulate cortex (vPCC, 10 bilateral), ventral medial prefrontal cortex (vmPFC, 10 bilateral), OFC (adjacent and more lateral to the color imagery OFC areas, 2 right, 7 bilateral); v) map imagery activated parahippocampal gyrus (parahippocampal place area, PPA, 1 left, 1 right, 6 bilateral), LOC (7 left, 3 bilateral), posterior parietal area (1 left, 9 bilateral), precuneus (10 bilateral), vPCC (10 bilateral).

**Fig 2.**
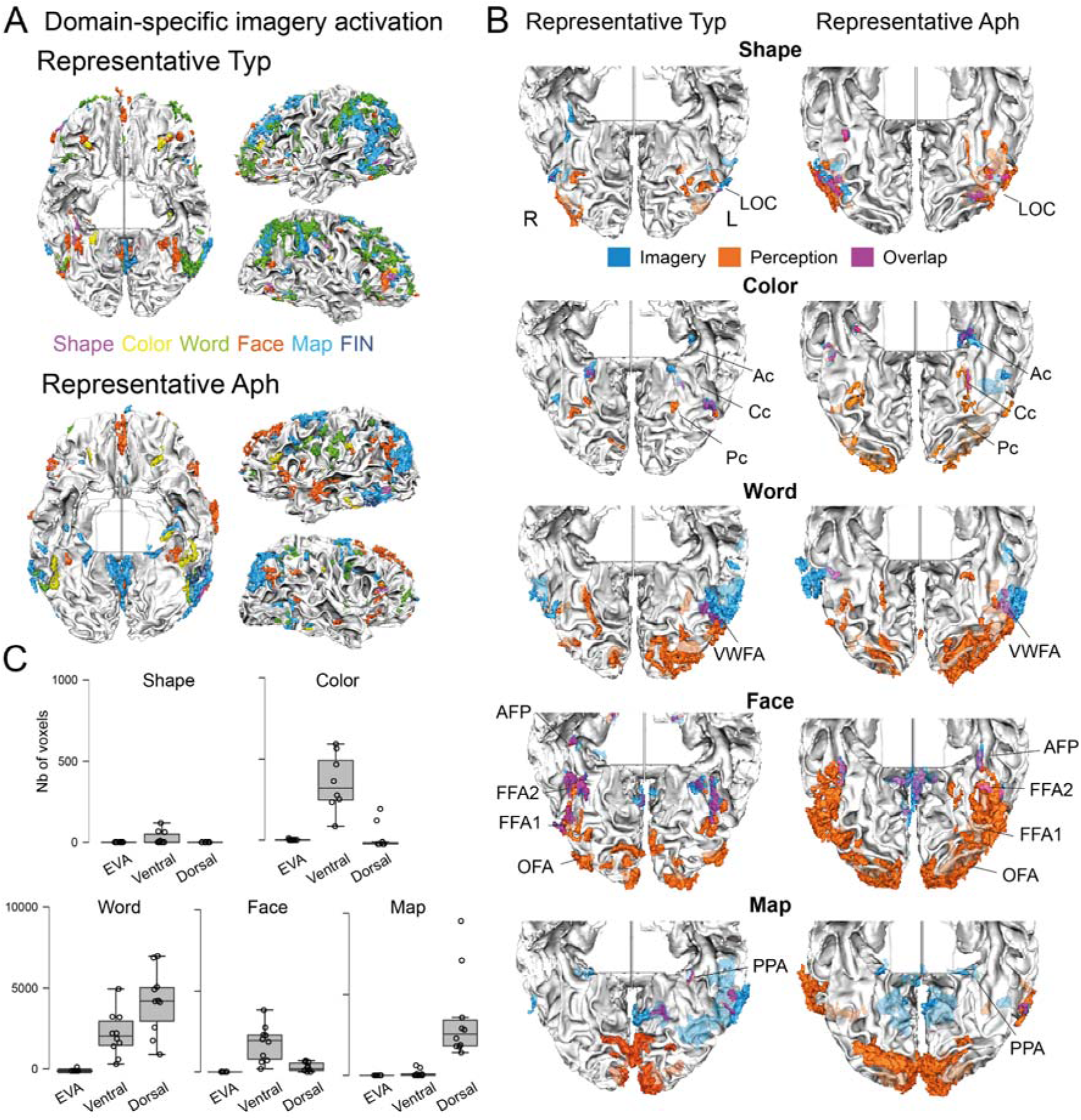
Patterns of BOLD response in domain-preferring regions during imagery, and their overlap with perception in typical imagers and aphantasic individuals. (A) Domain-specific activation during single imagery domains in a representative typical imager (Typ 05) and in a representative aphantasic individual (Aph10), from the contrast of one domain > the remaining 4 domains. All individual-level maps were thresholded at p<0.001, without any data smoothing, and cluster size > 12 voxels. The different domains and the FIN are color-coded. Note that aphantasic individuals showed clear domain-specific imagery activation in high-level visual areas. (B) Domain-specific activations during perception (orange), during imagery (blue), and their overlap (purple), in a representative typical imager (Typ 05) and in a representative aphantasic individual (Aph 10), unsmoothed data. Only the ventral view is displayed. Pc, Cc and Ac indicate the posterior, central, and anterior color-biased regions, respectively. Face patches include occipital face patch (OFA), fusiform face patches (FFA1 & FFA2) and anterior face patch (AFP). For all five domains, imagery-related activations overlapped with some of the perceptual-related activations in high-level VTC visual areas, but not with EVA despite stimulus-dependent EVA activation during perception. (C) In typical imagers, box-and-whisker plots of the number of voxels showing domain-specific activation during both mental imagery and perception, in EVA (V1, V2, and V3), and in the ventral and dorsal cortical visual pathways. Boxplot shows values of median, upper quartile, lower quartile, maximum and minimum, respectively. Dots represent single participants. Such unsmoothed voxels were present only in high-level visual areas, dependent on domain, but not in EVA.

**Table 1.**
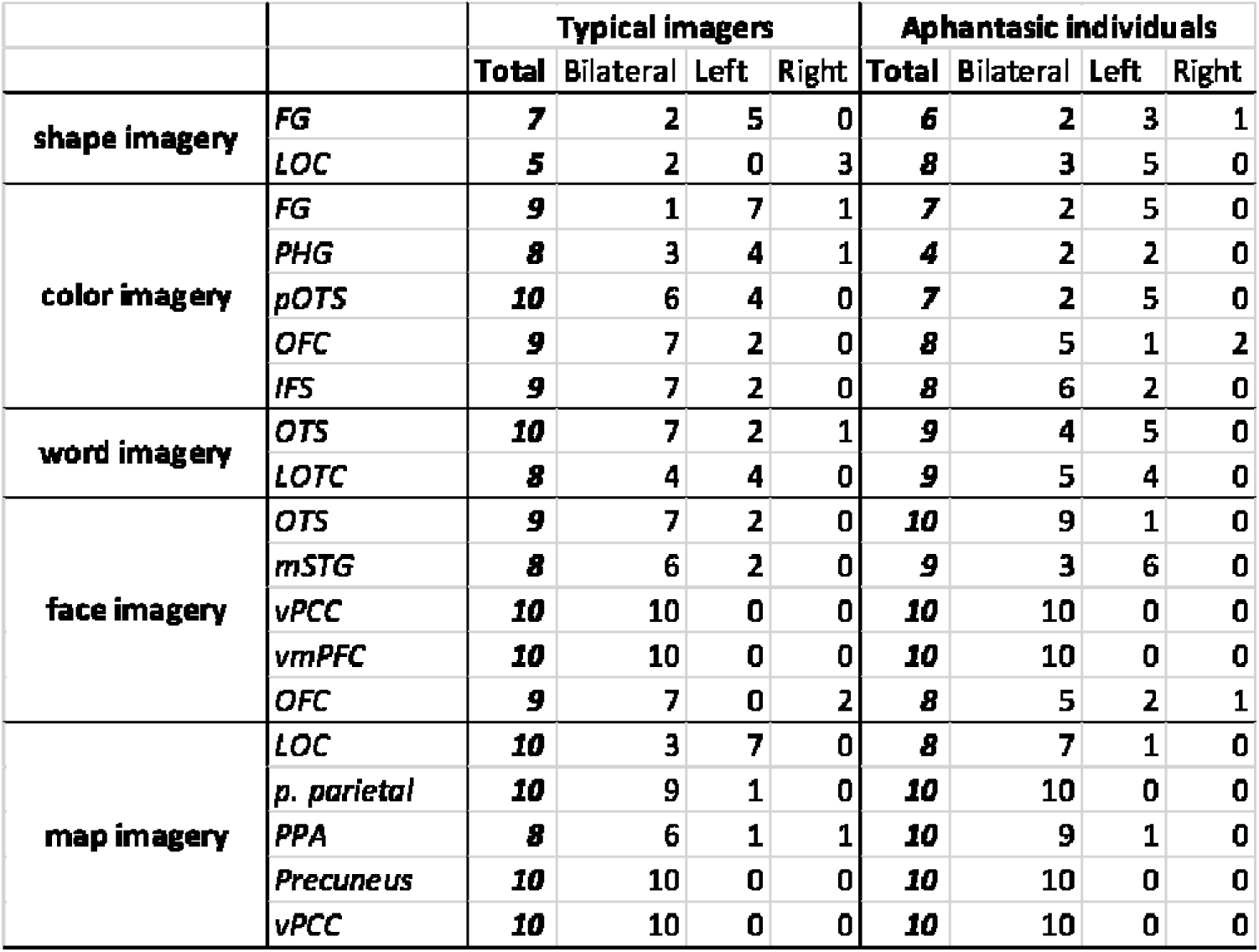
Number of participants exhibiting domain-specific activations during imagery.

See a list of glossary in supplementary materials. lFG: fusiform gyrus; IFS: inferior frontal sulcus; LOC: lateral occipital complex; LOTC: lateral occipitotemporal cortex; mSTG: middle superior temporal gyrus; PHG: parahippocampal gyrus; pOTS: posterior occipitotemporal sulcus; OFC: orbitofrontal cortex; vmPFC: ventral medial prefrontal cortex; vPCC: ventral posterior cingulate cortex; PPA: parahippocampal place area.

As expected, **perception tasks** evoked activity in both EVA and higher level visual areas (see Fig. 2B for a representative typical imager): i) shape perception activated the left LOC (Malach et al., 1995) and the bilateral medial FG (Kourtzi & Kanwisher, 2001); ii) color perception activated three patches in the bilateral medial FG, as well as the PHG (Lafer-Sousa et al., 2016) and the ventral OFC; iii) word perception activated the bilateral posterior FG (Cohen et al., 2000), and bilateral FP regions; iv) face perception activated the right OTS and bilateral amygdala (Kanwisher et al., 1997); v) the map of France activated the PPA, the bilateral vPCC, precuneus, and angular gyri, see (Epstein et al., 1999). Group-averaged results showing inter-individual consistency (if any) are displayed in Fig. S4 and Table S1 (for imagery) & S2 (for perception).

**Attempted imagery** in **<u>aphantasia</u>** evoked clear activations in domain-preferring regions (see Fig. 2A for a representative aphantasic participant, and Fig. S3 for the remaining participants; also see Table S5 for group-averaged results). Specifically, shape imagery activated the FG (3 left, 1 right, 2 bilateral out of 10 participants) and the LOC (5 left, 3 bilateral); color imagery activated FG (5 left, 2 bilateral), the pOTS (5 left, 2 bilateral), the OFC (1 left, 2 right, 5 bilateral), the IFS (2 left, 6 bilateral); word imagery activated the OTS (5 left, 4 bilateral) and the LOTC (4 left, 5 bilateral); face imagery activated the OTS (1 left, 9 bilateral), mSTG (6 left, 3 bilateral), vPCC (10 bilateral), vmPFC (10 bilateral), OFC (2 left, 1 right, 5 bilateral); map imagery activated PPA (1 left, 9 bilateral), LOC (1 left, 7 bilateral), posterior parietal cortex (10 bilateral), precuneus (10 bilateral), vPCC (10 bilateral). On **perception tasks**, aphantasic individuals activated similar domain-specific regions as typical imagers (Table S6).

### Comparison between aphantasic and typical participants

First, we compared **the extent of activation** between the two groups. We computed the number of activated voxels in domain-preferring areas, taking into account the individual variability in the location of activated regions. For each individual, we extracted the number of voxels in the volumetric intersection between domain-specific activations and EVA, the ventral visual pathway, and the dorsal visual pathway (see Methods for the specific areas), respectively. Individual domain-preferring patches consisted of unsmoothed voxels. We conducted a 3-way Bayesian ANOVA with the factor of Group x Region (including EVA and the ventral and dorsal cortical visual pathways) x Domain. There was moderate evidence supporting the absence of difference between aphantasic individuals and typical imagers, with no interaction with domains (Main Group effect, BF = 0.21 without interaction effects; see Fig.S5).

Second, we compared the **overlap between activations induced by imagery and by perception** between the two groups. When studying unsmoothed individual patches, for all five domains, there was some overlap between activations induced by imagery and by perception, in high-level VTC visual areas, in FP, and in subcortical regions (e.g. amygdala for faces). In contrast such overlap did not exist in EVA, despite domain-dependent EVA activation during perception (Fig. 2B). For example, color perception evoked activation in three color patches located bilaterally along the medial FG, the most anterior belonging to the VTC and the other two to EVA, as previously described by Lafer-Sousa et al. (2016).

During color imagery tasks, only the anterior or central color patches were activated, whereas the posterior color patch did not show any significant activation. In addition, the location of areas where imagery and perception overlap was domain-dependent; e.g. colors and faces engaged ventral patches whereas maps involved more dorsal patches (Fig. 2C).

In summary, typical imagers and individuals with aphantasia displayed similar imagery-related activity in the relevant domain-preferring areas, with some overlapping with perception-related activity in VTC patches.

### Univariate activation and multi-voxel pattern of the FIN

We then studied the activation of the FIN, a left VTC region which according to a meta-analysis (Spagna et al., 2021) is reproducibly activated during imagery. We compared the two groups using univariate measures, and multivariate methods probing representational format, in order to answer three questions: Do the current data support the hypothesis that a specific left VTC region is activated during imagery irrespective of content? Does this region also contain domain-specific information? Do aphantasic individuals differ from typical imagers in those respects?

**FIN activation amplitude. In typical imagers**, we identified domain-general regions by comparing all averaged imagery domains minus the abstract word task. This contrast showed a left-predominant set of regions (see Fig. 3A and Table S1) including the bilateral inferior frontal gyrus (IFG) and dorsal premotor cortex (PMd), the left intraparietal sulcus, and a region in the left posterior OTS (-41, -55, -10). This latter region was located at coordinates nearly identical to those of the FIN, as identified in our previous meta-analysis of fMRI studies of mental imagery (-40, -55, -11) (Spagna et al., 2021). **In aphantasia**, the same contrast showed activations in the left IFG-IPS network, right dorsal FP areas, and, importantly, also in the FIN (-45,-54,-10) (Fig. 3A and Table S5). How is the FIN activated during perception compared to imagery? If so, does activation differ between groups? We performed a ROI-based three-way Bayesian repeated measure ANOVA with the factors of Group x Modality [Imagery/Perception] x Domain. There was strong evidence for an absence of group differences of FIN activation amplitude (main Group effect, BF = 0.08; see Fig. 3A the activity profile of the FIN across tasks). FIN activation was stronger for perception compared to imagery (main Modality effects, BF = 194, partial eta-squared = 0.67) and was stronger for words than other domains (main Domain effect, BF = 5.65e5, partial eta-squared = 0.61). No interaction effect was found (all BFs < 0.14, all partial eta-squareds < 0.15).

**Fig 3.**
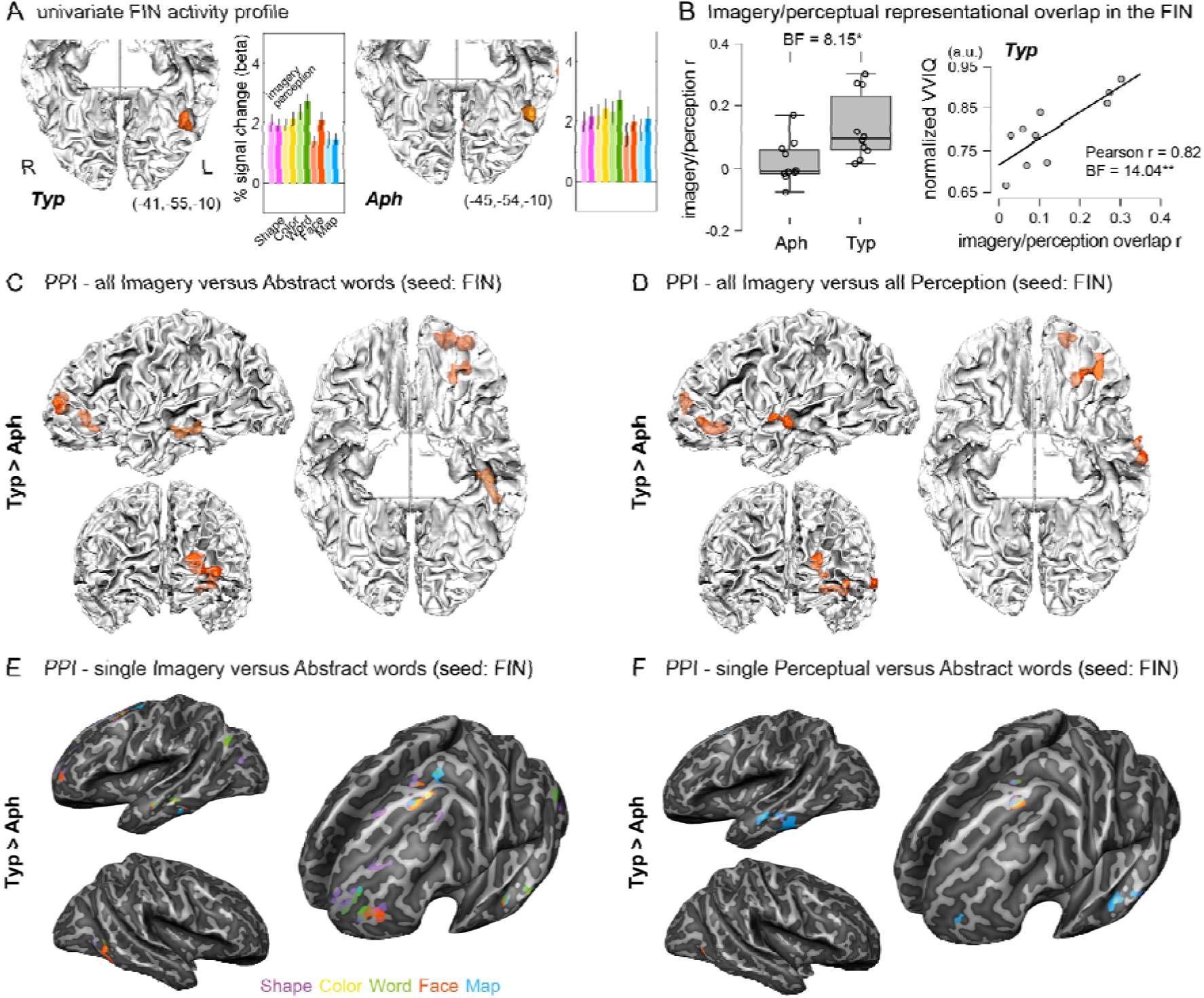
Activation profile, representational similarity and functional connectivity of the FIN between typical imagers and aphantasics. (A) Group-level contrast of all Imagery minus Abstract words identifies the FIN. Orange-colored patches indicate higher activation in imagery tasks than in the Abstract words task. All group-level maps underwent cluster-size thresholding using Monte-Carlo simulation with an alpha level of 0.05 and 5,000 simulations, and an initial threshold of p<0.005. The functional data was smoothed with a full width at half maximum of 6 mm. Group-level histogram of signal change in the FIN during imagery and perception for each domain. Error bars represent +/- 1 normalized SE across participants. Typ, typical imagers; Aph, aphantasic individuals. (B) Group difference of the correlation of imagery-perceptual RSMs. Dots represent individuals. Typ, typical imagers; Aph, aphantasic individuals. In typical imagers, correlation between individual Imagery-perceptual (I-P) similarity score r with subjective vividness (VVIQ score) in the FIN. VVIQ score was translated to a 0-1 scale. (C) Group difference in the functional connectivity of the FIN: all Imagery versus Abstract words. All group-level maps were thresholded at p<0.005 for cluster size correction. Orange shows regions with higher connectivity in typical imagers than in aphantasics. The FIN in typical imagers showed stronger connectivity during imagery with the left anterior PFC, OFC, and MTG/STG compared to aphantasic individuals. (D) Group difference of imagery domain-general FIN connectivity: all Imagery versus all Perception. Orange shows regions with higher connectivity in typical imagers than in aphantasics. (E) Group difference of single Imagery minus Abstract words FIN connectivity. In aphantasia, no FP region was significantly more connected with the FIN. The left PMd and anterior PFC regions were more connected to the FIN in typical than in aphantasic individuals across imagery domains. See 3D volume ROIs visualizations and detailed report for each domain in Fig. S8B. (F) Group difference of single Perception minus Abstract words FIN connectivity. In aphantasia, no FP region was significantly more connected with the FIN and no difference in connectivity was measurable during map perception. See also 3D volume ROIs visualizations and detailed report for each domain in Fig. S9A.

Does FIN activity actually depend on the anatomically close visual word form area (VWFA) activity? The FIN consisted of a single patch in all individual participants, unlike the domain-preferring regions, which were organized in multiple patches (Fig. S3). Its peak location was mesial, rostral and ventral to the VWFA (VWFA was defined by the contrast words perception versus other domains; Bayesian t-tests on coordinates, two-sided: X BF = 2.57, Y BF = 3.20, Z BF = 6.59, see Fig. S3 for individual maps), with partial overlap of VWFA activation in the lateral OTS. We specifically tested for differences in activation by identifying FIN-unique voxels, after excluding the voxels which overlapped with the VWFA. There was extreme evidence for different activation profiles between the FIN unique area and VWFA (BF = 2,629, see Fig. S6 for detailed statistics and activity profiles). These results confirm that the FIN is a functionally unique region, different from VWFA.

### The representational content of the FIN

We tested whether the FIN contains domain-related information by computing representational dissimilarity matrices (RDMs) between the multivoxel spatial activation patterns (Kriegeskorte, 2008). We computed the pairwise correlations between the spatial patterns of BOLD response elicited by the 90 imagery items and by the 30 perceptual items.

For imagery tasks, the RDMs of both groups featured small blocks around the diagonal in the FIN (Fig. S7, darker blue areas), suggesting the presence of domain-related information in the FIN. In addition, the RDMs showed high similarity between typical imagers and aphantasic individuals (r = 0.09; BF = 2.32e5). For perception tasks, there were similar diagonal domain blocks patterns in the RDMs of FIN (rs > 0.27; BFs > 1.19e6; n = 435), which showed highly similar representation between groups (r = 0.24, BF = 1.44e5).

However, the correlation between imagery and perceptual RDMs, or representational overlap, of the FIN (see RDMs in Fig. S7A), where participants imagined or perceived the same items, was higher for typical imagers than for aphantasic individuals (Bayesian t-test, BF = 8.15, Cohen’s d = 1.22; Fig. 3B). Moreover, the strength of this correlation was strongly correlated with individual vividness VVIQ scores only in typical imagers (r = 0.82, BF = 14.04, Fig. 3B), i.e. the greater the similarity in voxel patterns between imagery and perception of the same stimuli, the more vivid the mental imagery experienced by participants.

### Functional connectivity of the FIN

We examined the whole-brain functional connectivity of the FIN using psychophysiological interaction (PPI) to map its task-specific functional network during the imagery tasks and test whether aphantasia shows alteration of these connectivity within the imagery network.

For **domain-general connectivity**, we first studied typical imagers. We looked for regions whose correlation with the FIN would be higher during Imagery than during the Abstract words task, and found bilateral OFC regions (individual FINs as seed regions, Fig. S8A). We then compared the connectivity of the FIN between the “all Imagery” and the “all Perception” conditions. During imagery, we observed higher connectivity with the left OFC, mSTS and the right anterior temporal lobe and during perception, bilateral posterior occipitotemporal visual areas. At the individual level, we observed higher connectivity with the dlPFC for 9 out of 10 typical imagers (Fig. S10). The pattern of connectivity was very different in aphantasic individuals. No brain region displayed increased functional connectivity with the FIN in either the contrast of all Imagery vs. Abstract words or the contrast of all Imagery vs. all Perception. At the individual level, only 5 out of 10 aphantasic individuals showed higher connectivity with the dlPFC. Directly comparing patterns of FIN connectivity between the two groups confirmed that typical imagers displayed stronger connectivity during imagery than for either Abstract words (Fig. 3C, Table S10) or Perception (Fig. 3D) with the left anterior PFC, OFC, and MTG/STG compared to aphantasic individuals.

For **domain-specific connectivity**, we compared each imagery or perceptual domain minus Abstract words using PPI analysis. In typical imagers, we observed increased FIN connectivity with dorsolateral FP areas, and with the relevant domain-preferring VTC regions and subcortical areas in both domain-specific imagery and perceptual tasks (Imagery: Fig. 3E, detailed report of ROIs in Table S3 and Fig. S8 for each domain; Perception: Fig. 3F and Table S4). In contrast, in aphantasic individuals, when comparing single imagery or perceptual domains with Abstract words, we did not observe any increase in connectivity between the FIN and dorsal FP regions. Concerning specific imagery domains, in aphantasic individuals the FIN displayed higher local connections with FG in color imagery and right vPCC in map imagery (Fig. 3E and Table S7). No increased connectivity was found for the imagery of shapes, faces, and maps. For perceptual domains, the FIN showed only local higher connectivity with areas in the occipitotemporal region for the perception of shapes, colors, words, and faces (Fig. 3F and Table S7). Word perception induced also a higher FIN connection with right anterior PFC. No significant connectivity was observed in the perception of maps. Importantly, this absence of measurable functional connectivity pattern is consistent across all aphantasic individuals. A direct comparison between the two groups revealed that left PMd and anterior PFC were regions consistently more connected to the FIN in typical imagers than in aphantasic individuals during both imagery and perception.

In task-residual connectivity, we regressed out the task-related activity to obtain task residual data in imagery runs, and derive a proxy for resting-state functional connectivity. We observed FIN connectivity with the bilateral supplementary motor area, precuneus, V1, left middle frontal gyrus, FG and right IPS in typical imagers (Fig. S9C) and with bilateral V1 and right MFG in aphantasic individuals. No significant group difference was observed in task-residual connectivity.

In summary, the FIN showed comparable activation and representational content between typical imagers and aphantasic individuals. In typical imagers, the FIN is functionally connected with FP areas and with the relevant domain-preferring regions during both imagery and perception. Aphantasic individuals showed no measurable functional connectivity between the FIN and other regions in either imagery or perception.

### Univariate and multivoxel analyses and V1 functional connectivity

If EVA plays a crucial role in mental imagery, then aphantasia might be associated with abnormal EVA activity patterns. To investigate this possibility, we conducted the same analyses for V1 as we did for the FIN.

For **activation amplitude**, we compared the activity in foveal and peripheral V1 ROIs between the two groups using Bayesian repeated measures ANOVAs (group x task). As expected, the foveal V1 was activated during the perceptual tasks, with no group differences (BF = 0.53, Fig. 4A). In contrast, we observed deactivation in foveal V1 for the imagery tasks and for the abstract word task in both groups. This deactivation had a lower amplitude in aphantasic individuals than in typical imagers during the imagery tasks (BF = 7.41), and during the abstract word task (BF = 3.47). No difference between tasks and groups was found in the peripheral V1 (BF = 0.27).

**Fig 4.**
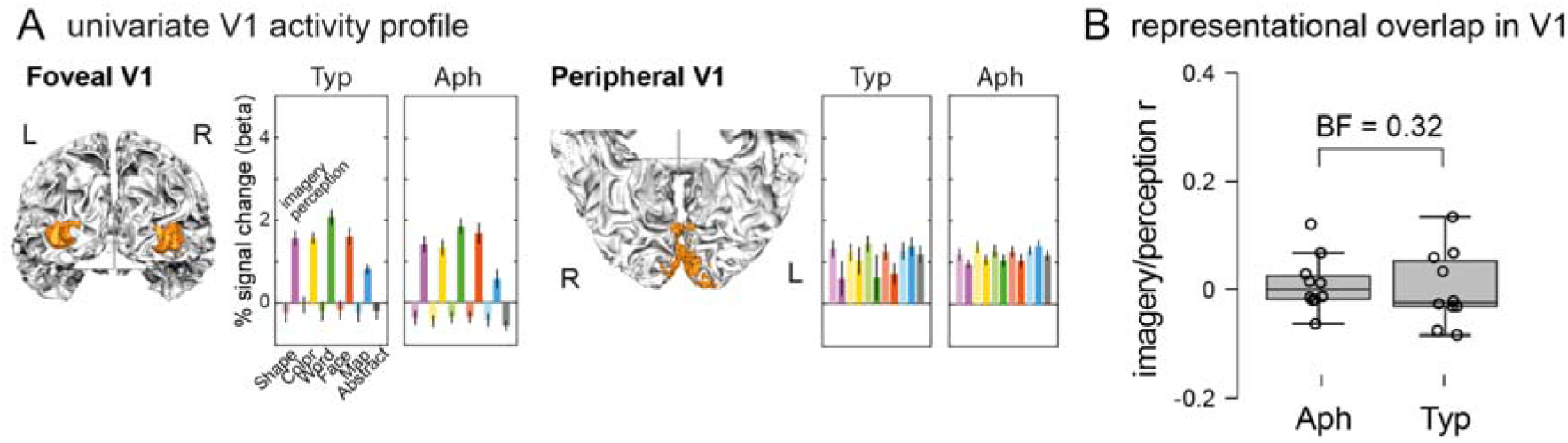
Activation profile, representational similarity and functional connectivity of V1 between typical imagers and aphantasics. (A) Group-level histogram of signal change in the foveal V1 and peripheral V1 ROIs during imagery and perception for each domain. Error bars represent +/- 1 normalized SE across participants. Light and dark colors refer to imagery and perception, respectively. Note the deactivation in foveal V1 during imagery and the abstract words task. Typ, typical imagers; Aph, aphantasic individuals. (B) Group difference of the correlation of imagery-perceptual RSMs. Dots represent individuals. Typ, typical imagers; Aph, aphantasic individuals.

For **representational content**, during imagery, there was no evidence of within-domain similarity across trials in V1 (rs<0.04, BFs<1.09), and no significant correlation of RDMs between groups (r = 0.05, BF = 3.24). In contrast, during perceptual tasks, RDMs for V1 showed very strong evidence of within-domain similarity blocks in both groups (rs > 0.39, BFs > 1.02e14), and very strong correlation between groups (r = 0.61, BF = 1.08e42). There was moderate evidence to reject the correlation between imagery and perception representation in V1 (BF = 0.24, Fig S7A) and difference between groups (BF = 0.32, Cohen’s d = 0.15).

Finally, we studied the functional connectivity of foveal and peripheral V1 seeds, and found no significant group differences with either task-specific or task-residual analyses (all ps>0.005, Fig. 4C).

### Higher activation of a right IFG-SMG network in aphantasia

Most previous analyses were focused on the VTC and EVA. We then assessed whether differences between groups existed also in non-visual brain sectors, particularly in the frontal and parietal regions with an involvement in mental imagery. We used whole-brain ANOVAs to compare brain activation. For the all Imagery tasks, aphantasic individuals showed higher activation than typical imagers in the right-hemisphere IFG, supramarginal gyrus (SMG), IPS, pMTG and a in a left occipital area (Fig 5A and Table S8). These same right-hemisphere FP regions were *deactivated* in typical imagers for the same contrast. Notably, during perception also, the same right-hemispheric IFG-SMG network showed higher activation in aphantasic individuals than in typical imagers (Fig. 5B). Activation during the control task with abstract words showed no activation difference between groups.

**Fig 5.**
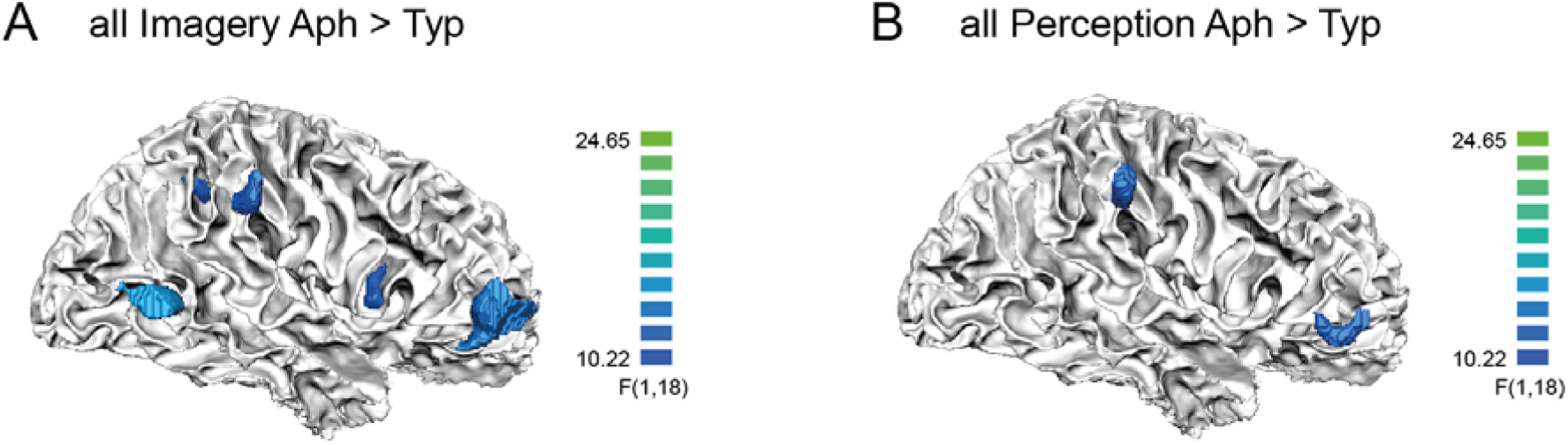
Higher activation in right-hemisphere regions in aphantasia. (A) Group difference in all Imagery tasks. Blue shows regions with higher activation in aphantasic individuals than in typical imagers. Typ: typical imagers. Aph: aphantasic individuals. (B) Group difference in all Perception tasks. Regions with higher activation in aphantasic individuals than in typical imagers are shown in blue.

Concerning domain-specific imagery activation, no areas other than the right hemisphere IFG-SMG network were differentially activated between groups.

Last, we sought to identify brain areas whose activity is modulated by subjective vividness across the imagery trials. While there was no significant group-level effect of vividness, the FIN appeared to be the most consistently modulated area by vividness when considering individual participants (4 out of 6 typical imagers who showed significant modulated regions, Fig. S11). No participant showed modulation of activity in EVA.

## Discussion

We used ultra-high-field fMRI to systematically examine domain-general and domain-specific mechanisms of visual mental imagery in typical imagers and in individuals with congenital aphantasia, who claim not to experience any visual mental imagery during wakefulness. Our study involved comprehensive testing of visual mental imagery capabilities across five different domains, namely object shapes, object colors, faces, letters, and spatial relationships. In both typical imagers and aphantasic individuals, imagery tasks activated the relevant domain-preferring VTC patches for each of the five explored imagery domains.

Importantly, imagery overlapped with perception only in the anterior VTC domain-preferring patches. In addition, we observed a domain-general cortical patch within the posterior lateral OTS in the left hemisphere, in a location consistent with the Fusiform Imagery Node (FIN) (Spagna et al., 2021). In aphantasic individuals, imagery and perception exhibited similar activation and representational content in high-level visual areas.

Beyond those commonalities, typical imagers and aphantasic participants differed in four respects, which may be helpful in understanding aphantasia. First, in the FIN, the imagery/perception overlap (ie, the correlation between imagery and perceptual representations) was greater for typical imagers than for aphantasic individuals. Only in typical imagers did this overlap correlate with subjective vividness measured by VVIQ. Second, in aphantasic individuals there was reduced functional connectivity between the FIN and frontoparietal areas. Third, aphantasic individuals showed enhanced deactivation of foveal V1 activity during imagery. Last, aphantasic individuals showed higher activation of a right-hemisphere IFG-SMG network. We will now discuss in turn the role of the VTC, the FIN, and the EVA in conscious imagery and in aphantasia.

### The role of domain-preferring ventral temporal areas

Dissociations in performance across various imagery domains among neurological patients (Bartolomeo, 2002; Goldenberg, 1993) suggest the existence of domain-specific circuits for visual mental imagery of object shape, object color, written words, faces, and spatial relationships, which may partly overlap with the corresponding domain-preferring circuits in visual recognition (Cohen et al., 2000; Epstein et al., 1999; Kanwisher et al., 1997; Lafer-Sousa et al., 2016; Malach et al., 1995). Our findings clarify the topographical organization of VTC regions thanks to high-resolution precision imaging in individual participants (see Fig. S3). As mentioned above, previous work had shown FFA and PPA activity during imagery of faces and places, respectively (Ishai et al., 2000; O’Craven & Kanwisher, 2000). We replicated those findings, and extended them by demonstrating imagery-related activity in the LOC for shape imagery, in the VWFA for letter imagery, and in color-biased regions for color imagery. Importantly, individual analyses allowed us to put to light the overlap of the domain- preferring patches activated during imagery and during perception. This overlap was restricted to the VTC high-level visual and associative areas. For instance, color imagery activated the anterior color-biased patches but not the more posterior ones.

Beyond the occipitotemporal cortex, we also found domain-preferring imagery activity in dorsal FP networks, in subcortical regions (such as in the amygdala for face imagery, possibly encoding face-associated emotions), and adjacent OFC patches for faces and colors, respectively. Face-preferring OFC patches had previously been described by Ref. (Barat et al., 2018; Tsao et al., 2008) in monkeys. The previously unknown color-preferring OFC patches may be related to the behavioral saliency of colors, particularly in detecting the emotional aspects of faces and food choices, such as assessing the ripeness of fruits (Siuda-Krzywicka et al., 2019).

### A domain-general imagery node in the left fusiform gyrus

The FIN was consistently active during visual mental imagery and during perception, independent of the imagery domain. Left temporal activity was previously described at locations close to the FIN during imagery (D’Esposito et al., 1997; Yomogida et al., 2004).

Anatomically, the FIN is contiguous to domain-preferring VTC regions such as VWFA or the FFA, such that one may ask whether it is topographically distinct from those regions. The present results localize the FIN around the left-hemispheric posterior OTS in 19 participants of 20. Individual analyses showed that the FIN, which was always restricted to a single patch, tended to be more mesial, rostral and ventral than the VWFA. Its location was sandwiched between the VWFA laterally and FFA mesially, with a possible partial overlap with these regions.

Our findings substantiate the hypothesis that the FIN has a very specific involvement in both imagery and perception, based on the following functional attributes. First, the FIN showed domain-generality by its increased BOLD activation when performing imagery tasks in all five domains: object shape, object color, written words, faces, and spatial relationships. Second, the FIN, together with the left-hemisphere IFG/IPS (see Fig. 6A), coded semantic content in its multivoxel patterns of activity. Third, the FIN showed increased functional connectivity with FP networks and with the relevant domain-preferring regions during domain-specific tasks, consistent with a role for the FIN as a semantic control hub (Jackson, 2021) for task-relevant features in both imagery and perception. Moreover, the strong left lateralization of the FIN, as well as that of domain-general FP areas, is in line with abundant evidence on hemispheric asymmetries of voluntary generation of visual mental imagery (Farah, 1984; Liu et al., 2022) and in discriminating imagery from perception (Koenig-Robert & Pearson, 2020). Such hemispheric asymmetry is consistent with the predominant left- lateralization of the semantic system (Binder et al., 2009; Fernandino et al., 2022), which provides a main input to voluntary visual mental imagery. Thus, these results support the notion of a semantic contribution of the FIN to the construction of mental images.

Still, the stronger activation of the FIN during imagery tasks than during abstract words processing might have an alternative explanation: the FIN could merely be a semantic region rostral to VWFA, specialized for concrete as opposed to abstract words. This possibility, however, would be in line with the very definition of concrete words, which are words linked to sensorimotor-based experience. As a consequence, word concreteness is highly correlated with word imageability (r=.971, Khanna & Cortese, 2021). Our imagery tasks allowed us to examine the domain-specific visual mental imagery (e.g. for faces, words, colors etc.), which elicited very localized activation in domain-specific high-level visual areas. The domain-specificity examined here is completely independent of the word- concreteness effect. Critically, the positive correlation we observed in typical imagers between VVIQ scores and the overlap in representation between imagery and perception in the FIN supports the notion that the eBIP trials successfully elicited imagery experiences in these participants. Thus, the activity of the FIN would not only result from the concreteness of the stimuli, but is related to experiencing the imagery itself. In other studies, the FIN appeared to be critically involved in mental imagery in studies using a variety of contrasts, some of which implemented non-verbal cues to evoke imagery (Spagna et al., 2021). For example, left fusiform activations similar to ours have been found in studies using non-verbal stimuli such as drawings (Mazard et al., 2005) or mathematical formulas (Pyke et al., 2017). In a further study (Amalric & Dehaene, 2016), mathematicians rated their subjective imageability of mathematical statements. These ratings positively correlated with activity in a left inferotemporal region (Talairach coordinates: -54; -52; -1) close to the FIN (Talairach coordinates: -41, -55, -10).

The separation between domain-general and domain-specific functions in the VTC is likely to minimize the cost of long-distance wiring (Sporns & Betzel, 2016) thanks to locally dense connections between domain-preferring regions and the FIN, and sparser connections with more remote areas. The FIN, equipped with long-range connections with the perisylvian language network and the anterior temporal lobe (Hajhajate et al., 2022), may thus act as a central hub for global back-and-forth communication between visual areas and language-related regions. Thus, the FIN may act as a bridge between semantic and visual information, enabling the generation of mental images with a visual content. The observation of functional connectivity between FIN and dlPFC / OFC dovetails nicely with the anatomical connectivity of the FIN to these regions, and supports its role as a domain-general node at the confluence of top-down influences from the FP networks and horizontal connections with the VTC domain-preferring regions.

This hypothesis also offers an explanation for the deficits in visual mental imagery observed in neurological patients, and yields testable predictions. Patients with lesions affecting the domain-general FIN, or disconnecting it from semantic networks, are likely to experience general imagery impairments (Bartolomeo, 2021; Moro et al., 2008). Lesions or disconnections that specifically target domain-preferring regions may result in more domain-specific patterns of mental imagery deficits (Bartolomeo et al., 2002). More posterior lesions disconnecting VTC from visual input are instead likely to produce perceptual deficits with preserved visual mental imagery (Bartolomeo, 2002, 2021).

### The relationship between visual perception and visual mental imagery

Our findings elucidate three important aspects of typical visual mental imagery, indicating that imagery does share some neural substrates with visual perception (Dijkstra et al., 2019; Mechelli et al., 2004), but with prominent differences. First, domain-preferring VTC regions exhibited some overlap between imagery and perception in more rostral patches, whereas more caudal patches only responded to perception. Second, we identified shared cortical patterns of representation for semantic domain content between imagery and perception in the high-level visual cortex. This finding aligns with previously reported similarities in domain-preferring visual areas between imagery and perception (Cichy et al., 2012; Reddy et al., 2010; Stokes et al., 2009). Notably, this representational overlap correlated with the level of vividness in typical imagers, specifying an objective measurement of subjective vividness across imagery domains, consistent with previous findings of representational overlap in high-level visual cortex (Dijkstra et al., 2019). Thus, the FIN may engage perceptual representations which allow to simulate vivid quasi-perceptual experience in imagery. Third, the FIN displayed different functional connectivity patterns between imagery and perception. In imagery, it displayed stronger connections with semantic networks, whereas in perception, it showed greater connectivity with occipitotemporal areas. Taken together, this evidence emphasizes the importance of high-level visual cortex for imagery and the common role of the FIN in processing semantic and visual content for both imagery and perception.

### The role of early visual areas in visual mental imagery

In both typical imagers and aphantasic participants, we observed peripheral V1 activation during imagery and perceptual tasks, as well as during the control task with abstract words. This activity might result from orienting of spatial attention in response to auditorily presented stimuli, which is often found in peripheral retinotopic locations of V1 (Brang et al., 2015; Cate et al., 2009), even in the absence of external stimuli (Kastner et al., 1999). Importantly, however, *foveal* V1 was active in perception but showed negative activity in imagery and in the abstract words task. This can result from attention being endogenously directed toward internal thoughts, which may inhibit foveal V1 to prevent potential interferences from external input. These findings challenge standard models stressing the role of EVA in visual mental imagery (Kosslyn et al., 2001; Pearson, 2019). However, the pattern we observed is quite consistent with extensive neuroimaging evidence in neurotypical individuals, which shows that visual mental imagery triggers activity in VTC and FP networks - but not in the EVA (Mechelli et al., 2004; Spagna et al., 2021). Moreover, detailed studies of neurological patients provided causal evidence through observations of disrupted imagery following left temporal damage rather than following lesions restricted to the occipital cortex (Bartolomeo, 2002; Bartolomeo et al., 2020; Liu et al., 2022). In different experimental contexts, mental imagery of colors (Bergmann et al., 2024), or the expectation to see gratings (Aitken et al., 2020), have been shown to modulate activity in the deep layers of V1. However, the comparable V1 activity and connectivity of aphantasia in our study suggests that these V1 patterns have no causal contribution to conscious imagery experience, consistent with early suggestions (Crick & Koch, 1995).

### Functional disconnection in aphantasia

Surprisingly, aphantasic individuals exhibited the activation of similar brain networks during mental imagery as observed in typical imagers. This was confirmed by univariate and RSA analyses of BOLD responses, in both domain-general and domain-preferring VTC areas, and in the FP networks. Importantly, aphantasic individuals could generate imagery-related representational patterns similar to those of typical imagers, indicating the presence of relevant visual information in the high- level visual cortex during attempted mental imagery. Consistent with this observation, aphantasic individuals were able to perform as accurately as typical imagers on tests of mental imagery (Liu & Bartolomeo, 2023). However, the representational overlap between imagery and perception was decreased in aphantasic individuals compared to typical imagers. This finding suggests reduced perceptual/imagery matching in aphantasia. In line with this possibility, the observed higher activity of SMG in aphantasia might correspond to a “mismatch” signal between representations (Doricchi et al., 2022). Aphantasia could be accompanied by atypical processing of internal states, such as emotion processing and interoception (Kvamme et al., 2024). These features may be associated with aphantasia but do not necessarily define it. We also observed enhanced deactivation of foveal V1 activity during imagery, which may reflect a failure in the modulatory mechanism that suppresses non-imagined content (Pace et al., 2023). This could be related to feedback inhibition of low- level visual areas during imagery.

During imagery, typical imagers exhibited a functional connection between the FIN and FP network activity, consistent with previous findings of increased coupling between frontal and high-level visual areas during imagery compared to perception (Mechelli et al., 2004). However, such correlations were reduced in aphantasic individuals. In partial agreement with our results, Milton et al. (2021) found reduced resting-state functional connectivity between PFC and the visual–occipital network in aphantasia. Reduced long- range connectivity is a hallmark of various neurodevelopmental disorders, perhaps including aphantasia (Sokolowski & Levine, 2023). Within the framework of the Global Neuronal Workspace hypothesis of conscious perception (Dehaene et al., 2006; Mashour et al., 2020), such a functional disconnection between FIN and FP networks could be interpreted as depriving visual mental imagery of its conscious experiential content. In other words, VTC activity by itself might be sufficient to access visual information, but not to experience it as a conscious perception-like state. In line with our findings, the Global Neuronal Workspace hypothesis underscores the importance of long-range loops between different cortical areas in sustaining and broadcasting neural information across the brain, with the dorsolateral PFC cortex playing an essential role (Dehaene et al., 2006; Mashour et al., 2020). Hence, aphantasia, characterized as a relatively "pure" deficit in conscious imagery experience, with a paradoxically preserved ability to perform imagery tasks, offers a compelling testing ground for theories of the brain mechanisms of conscious experience.

Aphantasic individuals exhibited impaired ability for object imagery but not for spatial imagery (Blazhenkova & Pechenkova, 2019), as shown by OSIQ. However, our fMRI result showed a consistent functional disconnection for spatial imagery as for the remaining imagery domains in aphantasic individuals. This discrepancy may be due to differences in the tasks. In the OSIQ, the Object part directly evaluates the subjective vividness of mental images, for example, "My images are very vivid and photographic." On the other hand, the Spatial part primarily pertains to spatial knowledge without explicitly requiring imagery experience, for instance, "In high school, I had less difficulty with geometry than with art.".

These spatial questions differ substantially from our Map of France task, which requires subjects to visualize a map and assess the spatial location of imagined cities. Aphantasic individuals performed the task accurately, but when rating the trial-by-trial vividness they reported almost no mental imagery at all.

### Increased activity in aphantasia

In both imagery and perception, the present group of aphantasic individuals exhibited greater activity in right-hemisphere IFG and SMG, which are important components of the right-lateralized network for reorienting attention (Bartolomeo & Seidel Malkinson, 2019; Corbetta et al., 2008) and its interaction with conscious perception (Liu et al., 2023). Such abnormal activity may play a role in disrupting the subjective experience of generating or maintaining mental imagery, for example by interrupting ongoing activity in more dorsal FP networks (Corbetta et al., 2008). A possible mechanism could be defective filtering of distracting events (Shulman et al., 2007), leading to interference with internally generated models (Bartolomeo & Seidel Malkinson, 2022).

Several considerations support the notion that the imagery tasks of the eBIP were capable of evoking visual mental images. First, using the same battery, Liu & Bartolomeo (2023) found an inverse correlation between trial-by-trial subjective vividness and response times on the eBIP. Higher levels of vividness were associated with faster response times. Second, and more importantly, the eBIP imagery tasks induced activations of perceptual domain-preferring VTC patches, confirming and extending early findings by O’Craven & Kanwisher (O’Craven & Kanwisher, 2000) on imagery of faces and places. We obtained more systematic evidence here across the five semantic domains we investigated. Third, tasks similar or identical to the eBIP have often been used in clinical settings to assess domain-selective impairments. Neurological patients with imagery deficits in specific domains showed impaired performance on these tasks, which cannot be attributed to impaired semantic knowledge. For example, patient VSB (Bartolomeo et al., 2002) had a deficit in visual perception and visual imagery of letters (as assessed by questions on the visual shape of letters, similar to those used in the eBIP) as a consequence of a left temporal stroke. However, he could still answer the same questions when he was allowed to mimic their writing, demonstrating preserved semantic knowledge of letters.

Limitations of the present study include (1) The exclusion of bilateral anterior temporal lobes due to limited brain coverage at 7T (Fig. S2A). (2) The impossibility to analyze our trial-by-trial vividness scores, because of insufficient variability in the vividness ratings of typical imagers (consistently high) and of aphantasic individuals (consistently low; Fig. S1B). (3) We used the contrast of all averaged imagery domains minus the abstract word to identify the FIN. This contrast may introduce an additional concreteness effect beyond the imagery effect in the current data. Future studies may investigate whether the FIN would be involved in imagery of non-verbal stimuli to rule out this possibility. (4) The possibility that aphantasia might be a heterogeneous condition, in the absence of diagnostic criteria that could identify potential subtypes with differing neural substrates.

Despite these limitations, our findings shed light on the left-predominant circuits of individual-level visual mental imagery, encompassing the FIN, FP networks, and, importantly, domain-preferring VTC regions. This evidence suggests the presence of distinct domain-general and domain-preferring cortical components of visual mental imagery(Spagna et al., 2024). Our results also demonstrate that visual mental imagery and visual perception share similar neural mechanisms in the high-level visual cortex. Finally, we identified a neural signature of aphantasia, which was associated with reduced functional connectivity within the imagery network between the FIN and FP networks, despite essentially normal behavioral performance, BOLD activity levels, and representational content. Thus, the present results support the general hypothesis that conscious visual experience - whether perceived or imagined - depends on the integrated activity of FP networks and high-level visual cortex (Dehaene et al., 2006; Liu et al., 2023; Mashour et al., 2020; van Vugt et al., 2018).

## Acknowledgements

The research was supported by INSERM, CEA and specific funding from Dassault Systèmes. We are grateful to NeuroSpin support staff, and particularly to Bernadette Martins, the nurses and MR technicians at NeuroSpin who provided crucial help at various stages of data acquisition and analysis. The work of P.B. is supported by the Agence Nationale de la Recherche through ANR-16-CE37-0005 and ANR-10-IAIHU-06, and by the Fondation pour la Recherche sur les AVC through FR-AVC-017. J.L. expresses his gratitude for the 2023 European Workshop on Cognitive Neuropsychology Prize that was awarded to him in recognition of the work presented in this study.

## Contributions

Conceptualization: J.L., P.B., L.C., S.D.; Data curation: J.L., D.H., M.Z.; Formal analysis: J.L., M.Z.; Investigation: J.L., D.H., M.Z.; Methodology: J.L., M.Z.; Funding acquisition: P.B., J.L.; Project administration: P.B.; Resources: S.D.; Software: J.L., M.Z.; Supervision: P.B.; Visualization: J.L.; Writing - original draft: J.L.; Writing - review & editing: All authors

## Materials & Correspondence

Requests for materials and correspondence should be addressed to J.L.

