## Supplementary material for "Visual mental imagery in typical imagers and in aphantasia: A millimeter-scale 7-T fMRI study": V5

This PDF file includes:

Glossary list

Supplementary Figures 1 to 11

Supplementary Table 1 to 11

### Glossary list

| | | | |
|---|---|---|---|
| AC | anterior color | OFC | orbitofrontal cortex |
| AFP | anterior face patch | OTS | occipitotemporal sulcus |
| Aph | aphantasic individual | PC | posterior color |
| BF | bayes factor | PFC | prefrontal cortex |
| CC | central color | PHG | parahippocampal gyrus |
| dlPFC | dorsolateral prefrontal cortex | PMd | dorsal premotor cortex |
| eBIP | enhanced batterie imagerie / perception | PPA | parahippocampal place area |
| EVA | early visual areas | PPA | parahippocampal place area |
| FFA | fusiform face area | PPI | psychophysiological interaction |
| FG | fusiform gyrus | RDMs | representational dissimilarity matrices |
| FIN | fusiform imagery node | SMG | supramarginal gyrus |
| FP | frontoparietal | Typ | typical imager |
| IFG | inferior frontal gyrus | vmPFC | ventral medial prefrontal cortex |
| IPS | intraparietal sulcus | vPCC | ventral posterior cingulate cortex |
| LOTC | lateral occipitotemporal cortex | VTC | ventral temporal cortex |
| mSTG | superior temporal gyrus | VWFA | visual word form area |
| OFA | occipital face patch | | |

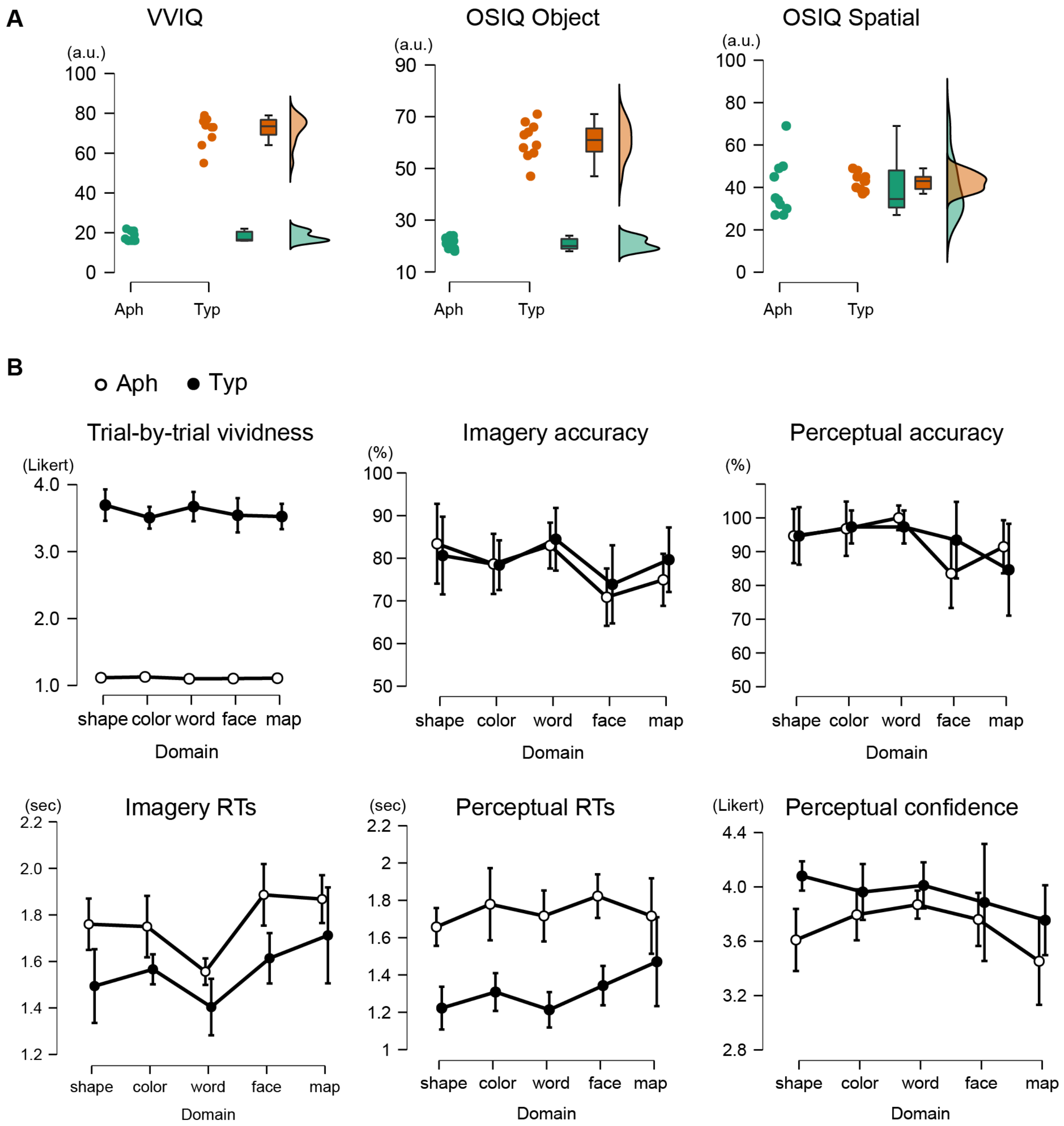


**Fig S1. Introspective reports and task performance.**
(A) VVIQ questionnaire scores and OSIQ questionnaire scores for object imagery and spatial imagery. Aph: aphantasic individuals. Typ: typical imagers.
(B) Task performance for aphantasic individuals and for typical imagers in imagery and perception tasks. a.u.: arbitrary unit. Trial-by-trial vividness and perceptual confidence rated on a Likert scale of 1 to 4. Error bars represent +/- 1 SD across participants. For imagery accuracy, there was no evidence for an Domain x Group interaction effect (BF = 0.11) but moderate

evidence for a difference between domains (main Domain effect, BF = 4.22). Post-hoc tests revealed lower accuracy in the imagery of faces compared to the imagery of words (BF = 48.69) or shapes (BF = 5.27). A similar domain difference was also observed in perception (main Domain effect, BF = 3.32; interaction effect, BF = 0.31), where the accuracy of face perception was lower than the accuracy of word perception (post-hoc tests, BF = 3.50).

**A**

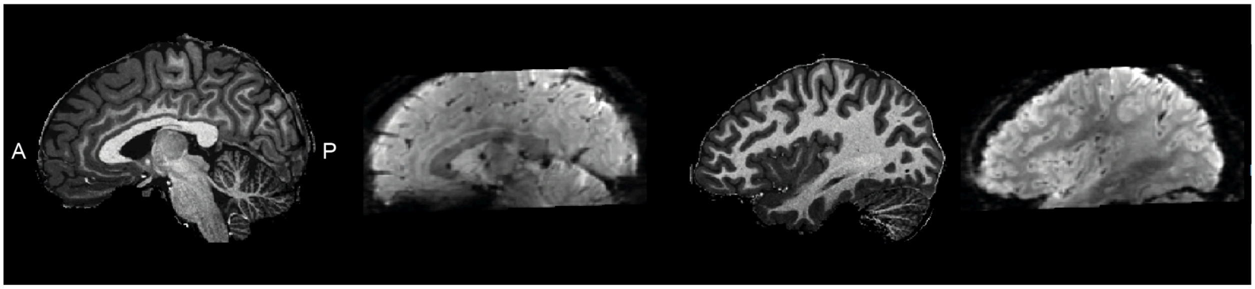


**B**

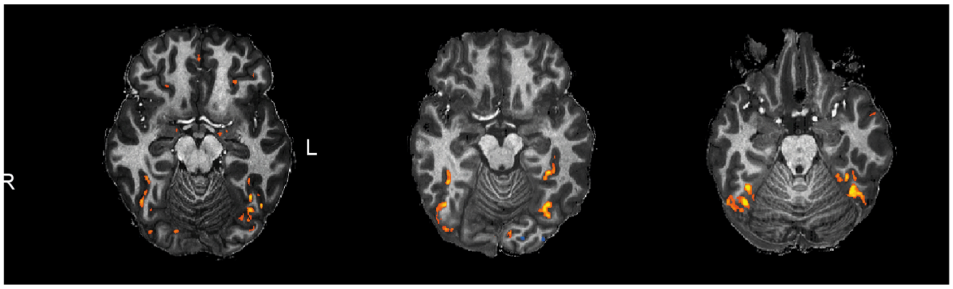


**Fig. S2 Brain coverage and Inter-individual anatomical/functional variability.**

(A) Sagittal view of T1 anatomical image (0.65mm isotropic, resampled to 0.6 mm isotropic) and functional image (1.20mm isotropic) in a representative participant. Part of the anterior temporal lobe was not able to be included.

(B) Face-specific activation in multiple ventral temporal cortex patches during face perception in three typical imagers, showing critical variability of the location of activated functional patches which reduced the overlap among participants in group-level analysis.

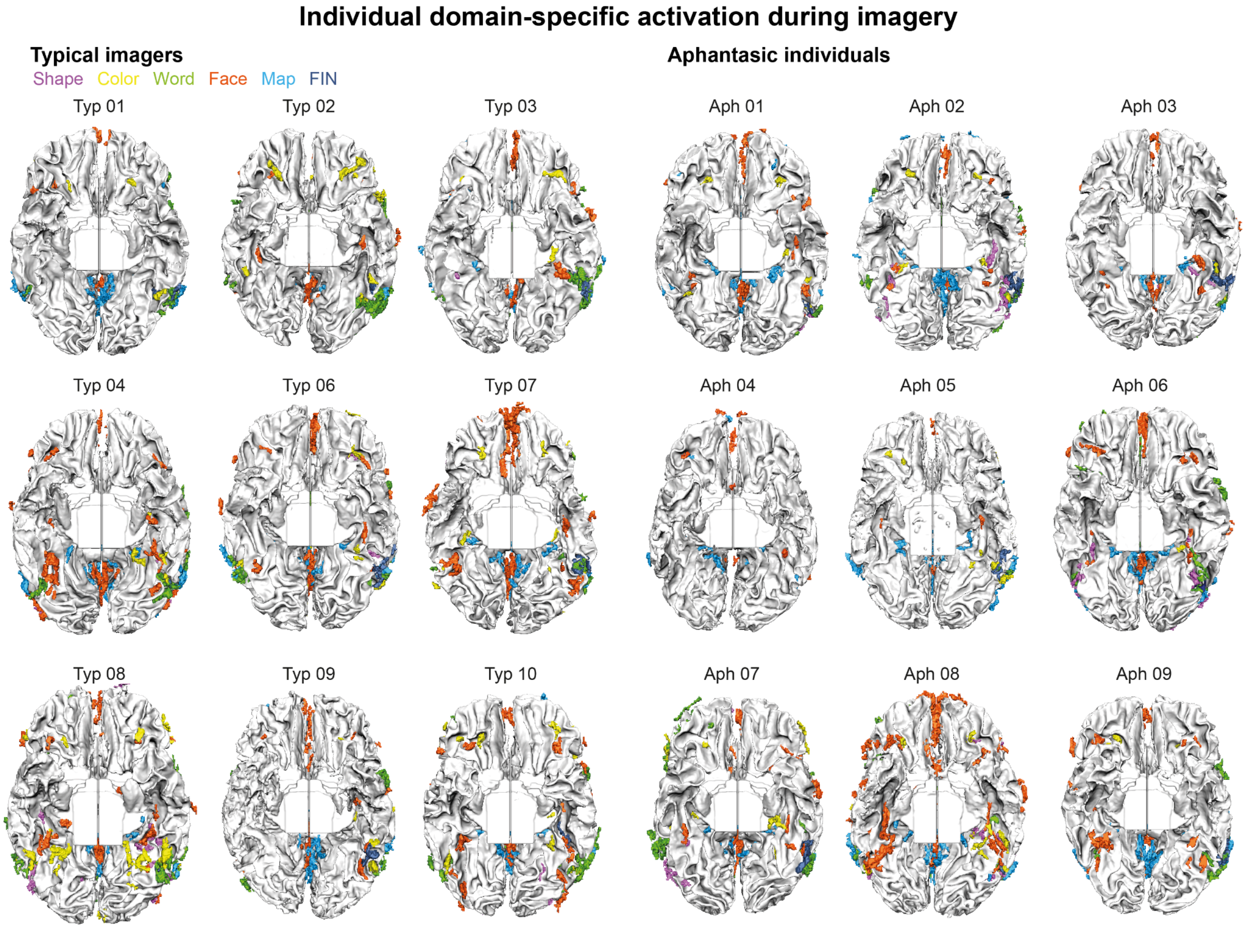


**Fig.S3. Individual domain-specific activations for imagery domains.** The domain of activation is indicated by color-coded words that use the same color. Typ.: typical imager, Aph.: aphantasic individual.

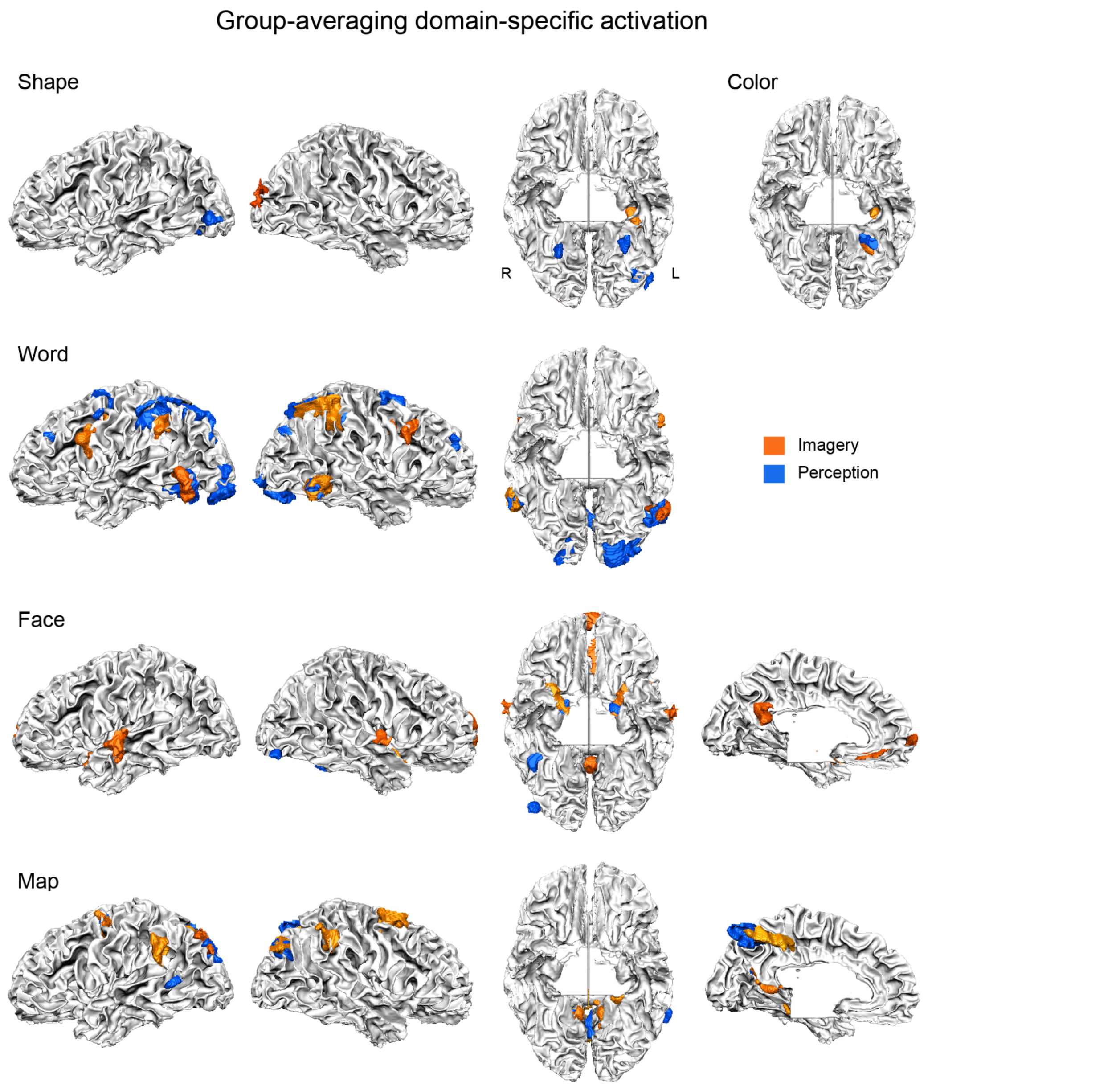


**Fig. S4. Group-averaging domain-specific activations during imagery and perception for typical imagers.** Group-averaging domain-specific activation that is greater than the activation in the other four domains during imagery domains (in orange) and during perceptual domains (in blue), showing consistency of activated location across individuals.

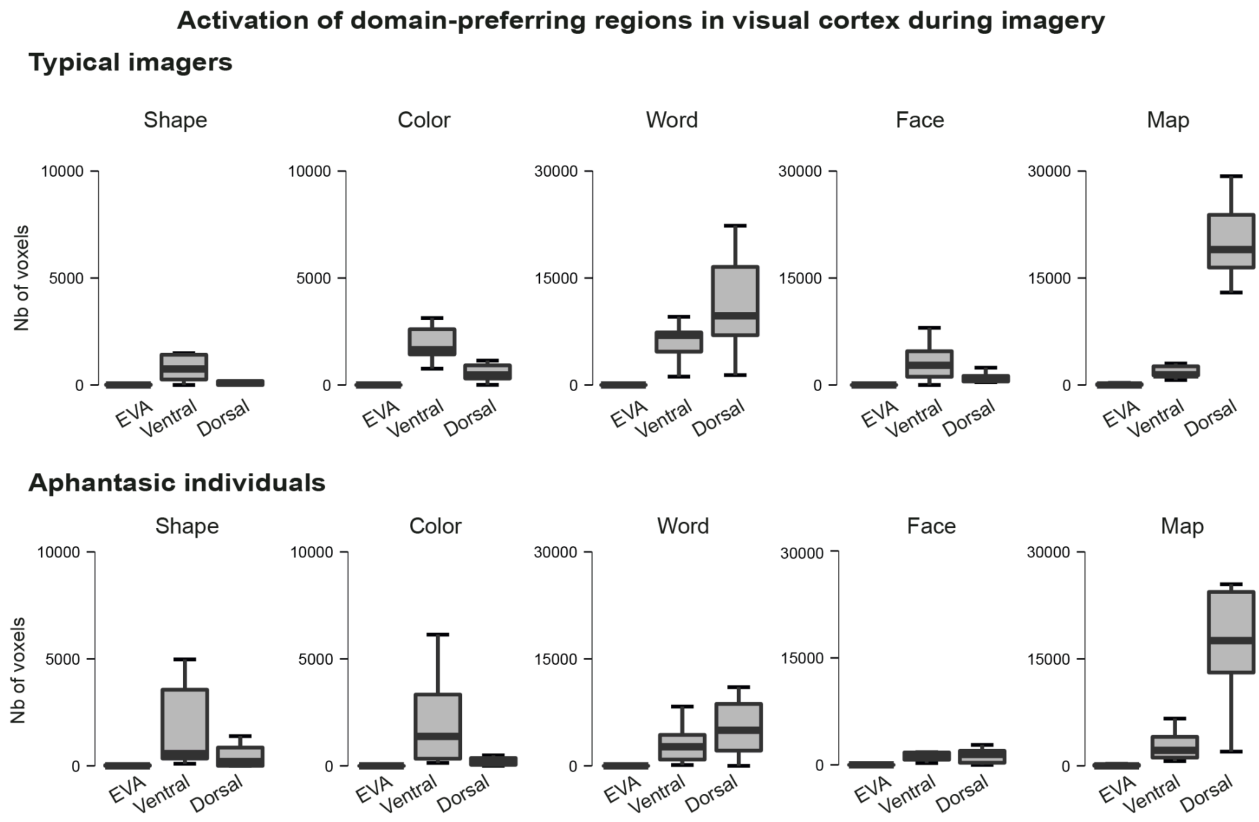


**Fig S5. Domain-specific activation in visual cortex during imagery.**
Number of voxels of domain-preferring regions activated in three visual regions (EVA, ventral and dorsal visual pathways) in each imagery domain. In the 3-way bayesian ANOVA with the factor of Group x Region x Domain. There were main Region and Domain effects (both BFs > 1.01e14) and a Region x Domain interaction (BF = 6.10e14). However, there was no evidence for Group x Domain x Region, Group x Domain, or Group x Region interactions (all BFs < 0.19).

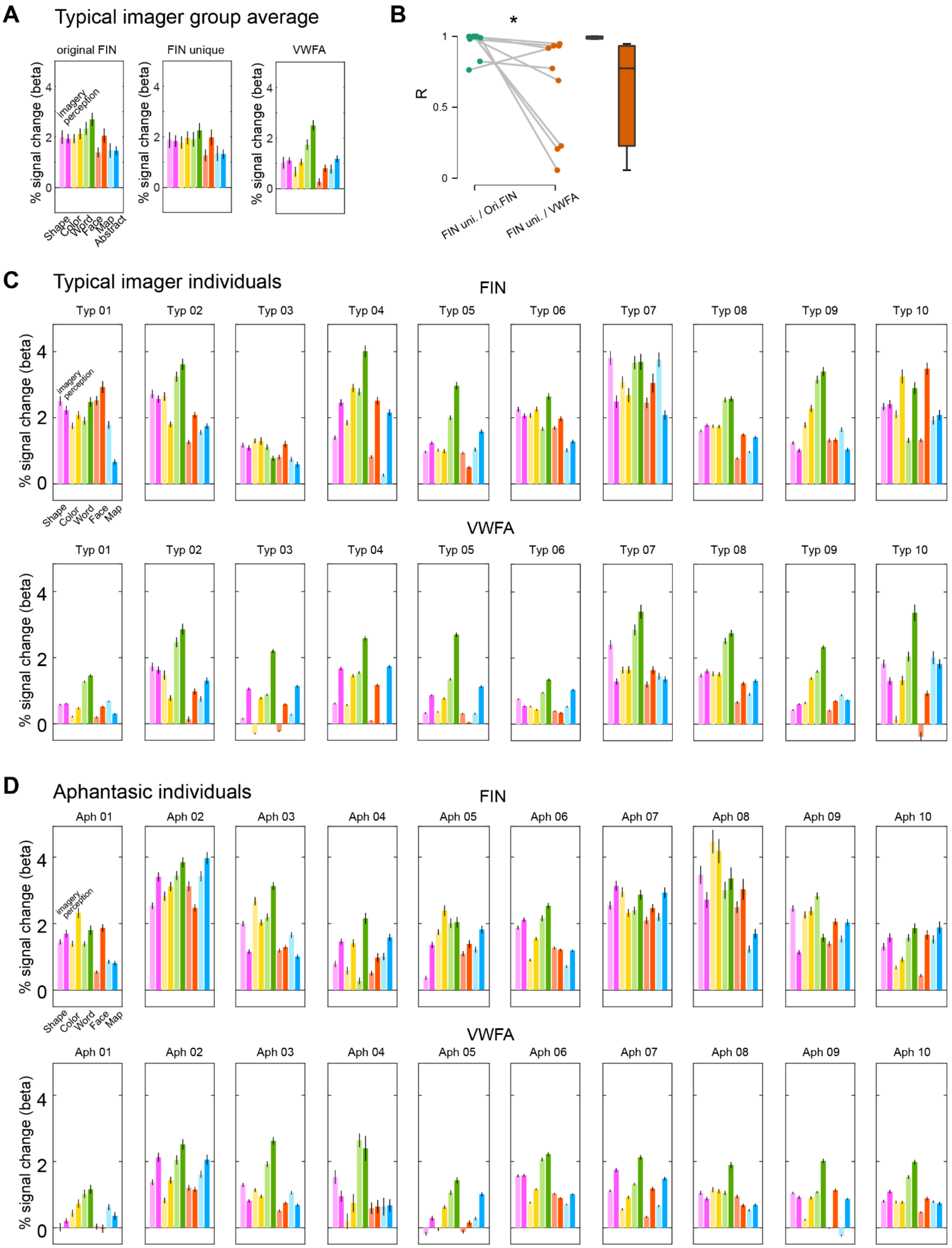


**Fig.S6. FIN and VWFA activation across domains of imagery and perception.** We examined the relationship between the FIN and the VWFA which was defined by the contrast of Word perception versus other perceptual domains. Although there may be some anatomical overlap in the left lateral OTS, the two functional regions showed different profiles of activation during both imagery and perception domains (see Fig.S6A for group-level average and Fig.S6C for typical imager individuals). The FIN was activated for each domain in imagery and in perception compared to baseline activities (beta value, Bayesian t-test, all BFs > 45.56, Fig.5E). In typical imagers, a Bayesian repeated measures ANOVA with the factor of ROIs (FIN, VWFA) and Task (imagery and perceptual domains) indicated very strong evidence for the different activation profiles between FIN and VWFA (main Region effect, BF = 68.11). Furthermore, to test possible biases in the FIN, we isolated the voxels that are unique to FIN (‘FIN unique voxels’) by excluding those that overlapped with the VWFA and performed the same analyses. There was no evidence for a difference of activation between the original FIN and FIN unique voxels (BF = 0.36 for the factor of Region). However, there was extreme evidence for the different activation profiles between the FIN unique area and VWFA (main ROIs effect, BF = 2629), confirming that the FIN as functionally unique region involved in the domain-general imagery. In addition, at individual level, the correlation between the activity profile of the FIN unique and the original FIN is higher than the FIN unique and the VWFA (Bayesian Paired Sample t-test, BF = 9.40). In aphantasia, we observed a consistent difference of activation profiles between the original FIN and VWFA (main Region effect, BF = 7.96).

(A) Group-level histogram of signal change in the original FIN, the FIN unique voxels and the VWFA during imagery and perception for each domain. The VWFA was defined by contrasting the perception of Words minus the perception of other categories. Error bars represent +/- 1 normalized SE across participants.

(B) Correlation of activity profile between the FIN unique (FIN uni.) voxels and the original FIN (ori. FIN), and between the FIN unique voxels and the VWFA. Dots represent individuals. * for BF>3.

(A) Signal change in typical imager individuals. Error bars represent +/- 1 SD across participants.

(B) Signal change in aphantasic individuals.

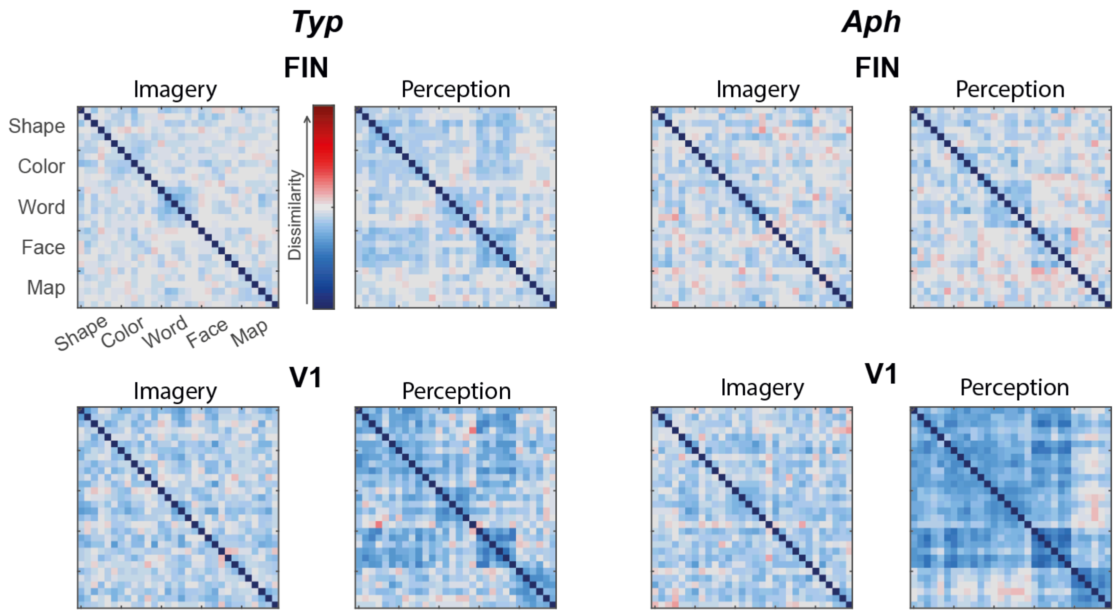


**Fig S7. Representational dissimilarity matrices in the same run.** Group-averaged representational dissimilarity matrix for imagery and perception of the same 30 items in the same run.

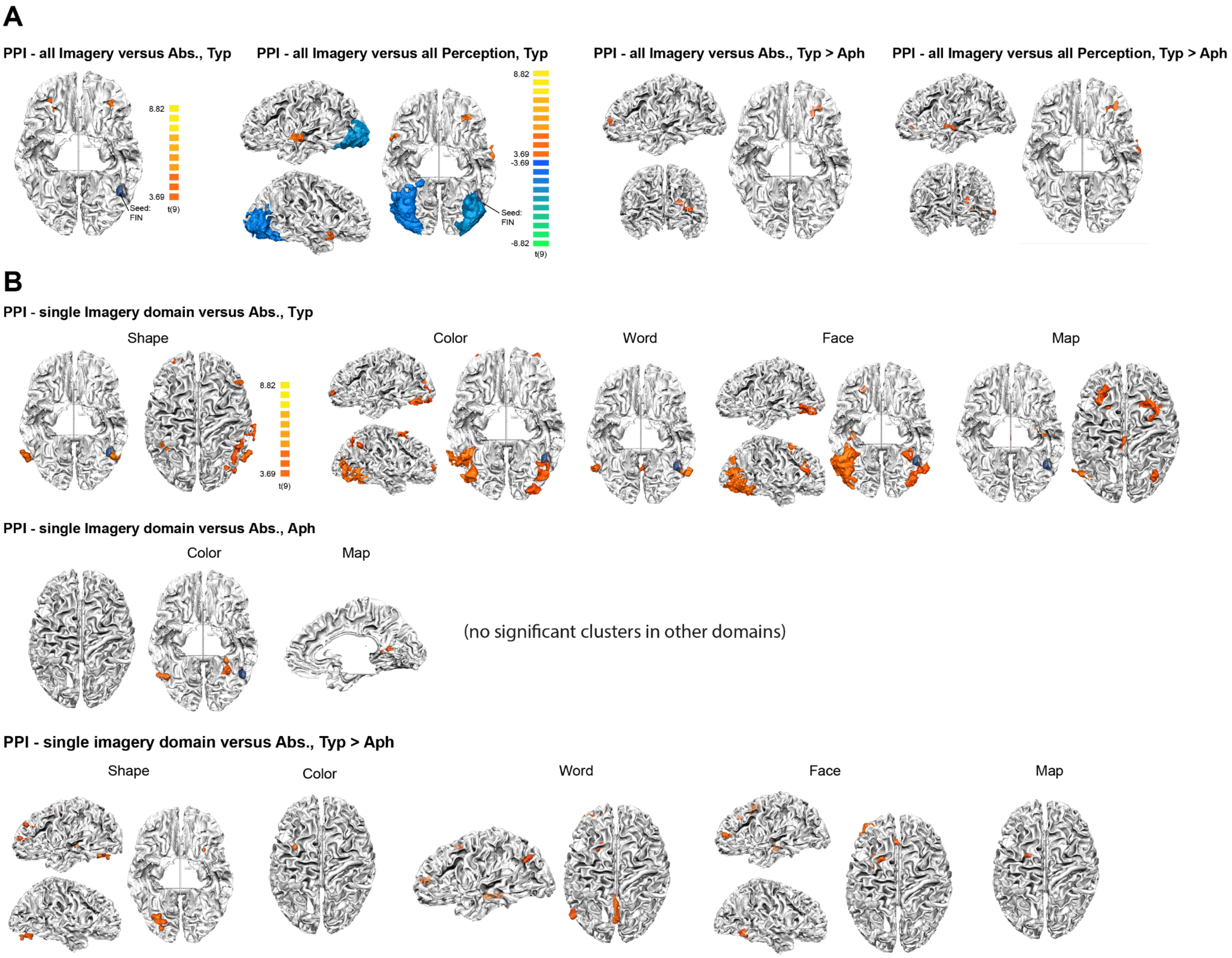


**Fig S8. Task-specific domain-general and domain-specific FIN connectivity during imagery.**

(A) PPI for domain-general FIN connectivity in typical imagers and group differences between typical imagers and aphantasic individuals. No significant cluster was found in aphantasic individuals for imagery. Typ.: typical imagers. Aph.: aphantasic individuals. Abs.:Abstract words.

(B) PPI for single Imagery domain *versus* Abstract words. In typical imagers (Typ), the FIN was connected with: bilateral lateral occipito-temporal areas, left IPSa, frontal pole, and right SMG, IPSa, MFG, for shape imagery; bilateral ANG, left MTG, SFG, right FG, SMG, IFG and OFC, for color imagery; bilateral fusiform gyrus and precuneus, for word imagery; bilateral amygdala, PMd, left MTG, PHG, vPCC, IPS and right FG, ANG, dmPFC, OFC, for face imagery; left PHG, ANG, SFS, caudate and right dPCC, IPS, PMd, for map imagery. For group difference (Typ > Aph), typical imagers showed higher FIN connectivity than aphantasic individuals (Aph): 1) in shape imagery, with left PMd, anterior PFC, dlPFC, Insula, MTG, and right lingual gyrus; 2) in color imagery, with left

PMd and MTG; 3) in word imagery, with left PMd, anterior PFC, ANG, MTG and bilateral precuneus; 4) in face imagery, with left PMd, FEF, anterior PFC, MTG and right ITG; 5) in map imagery, with left PMd and MTG.

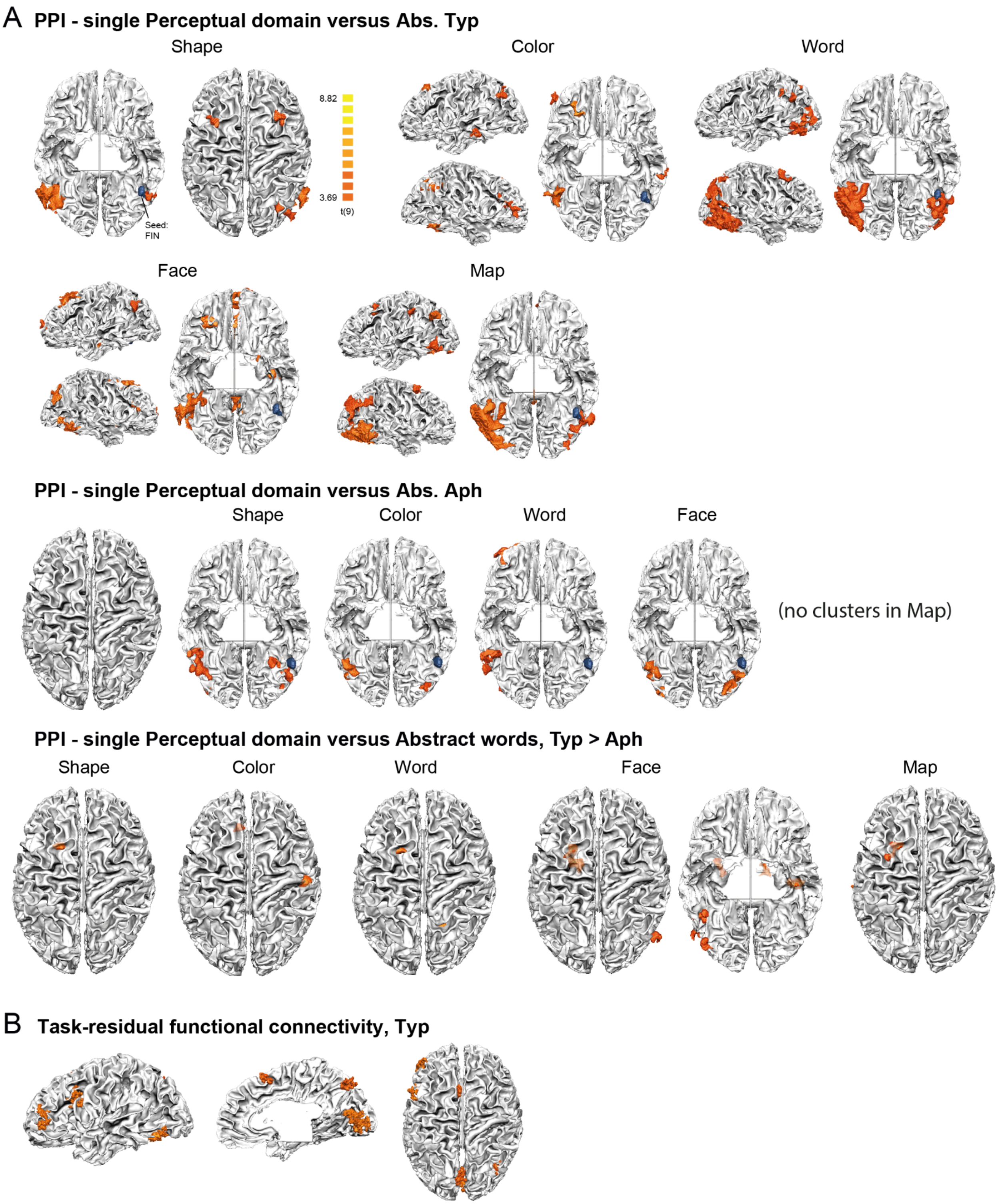


**Fig S9. Domain-specific FIN connectivity during perception and task-residual functional connectivity of the FIN.**

(A) PPI for single Perceptual domain *versus* Abstract words. Typ.: typical imagers. Aph.: aphantasic individuals. Abs.: Abstract words. Group difference of task-specific functional connectivity of the FIN in perceptual domains (Table. S11): typical imagers had higher

FIN connectivity: 1) in shape perception, with left PMd; 2) in color perception, with left FEF, MTG and right primary motor area; 3) in word perception, with left PMd, STS, right OTS and SPL; 4) in face perception, with left PMd, MTG, right LOTC, OTS and bilateral amygdala; 5) in map perception, with left PMd and MTG.

(B) Task-residual functional connectivity of the FIN. Talairach coordinates: bilateral supplementary motor area (-4, 14, 48), precuneus (-2, -67, 43), V1(1, -76, 7), left MFG (anterior: -43, 45, 6; posterior: -47, 14, 26), FG (-49, -61, -7) and right IPS (34, -59, 41). No group difference was found between typical imagers and aphantasic individuals.

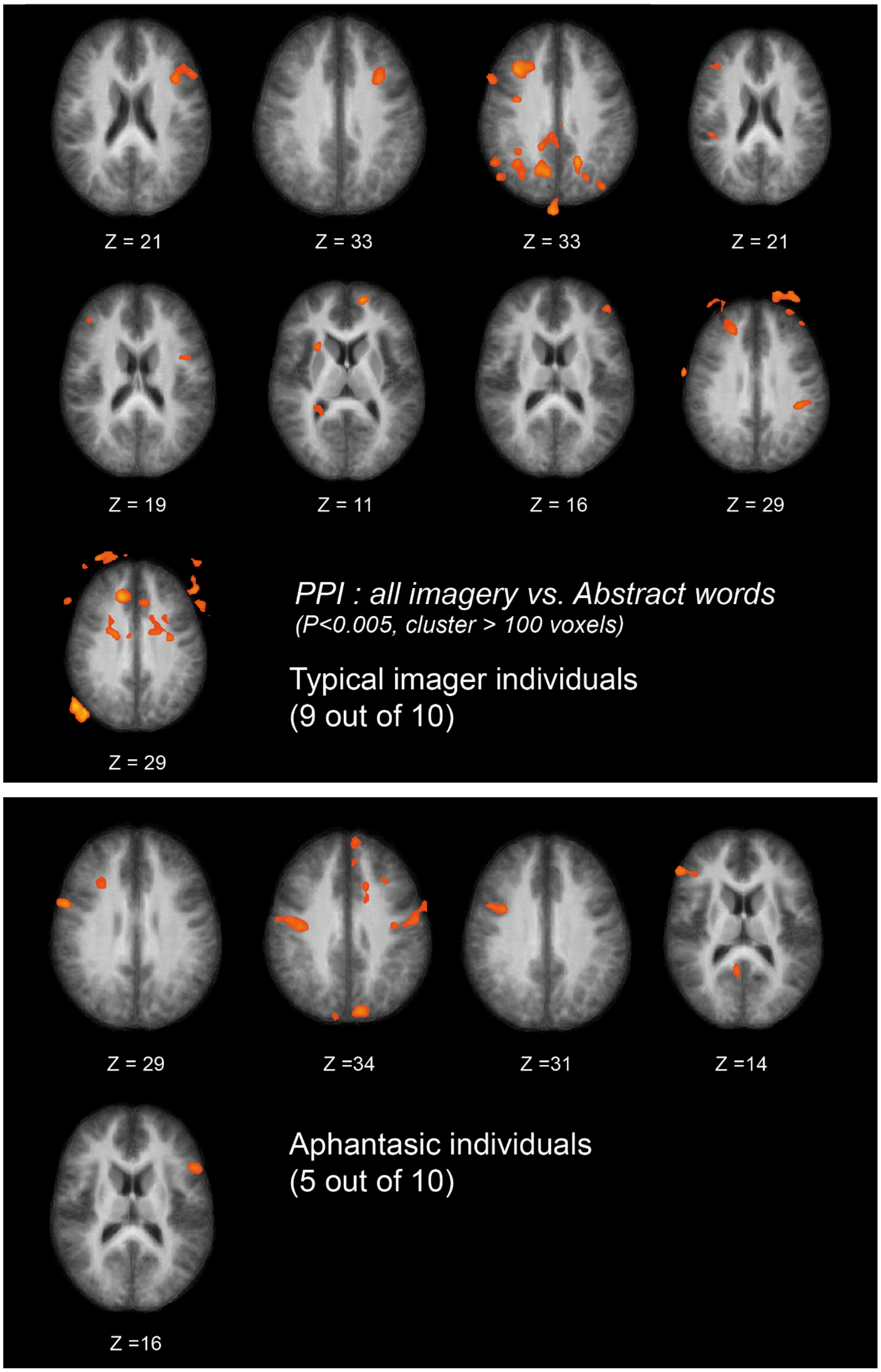


**Fig S10. Individual PPI functional connectivity in the contrast of all imagery versus Abstract words control condition** The seed was the FIN. Brain activation of each individual was plotted on the group-averaged brain anatomy. Each brain represents one participant. Nine out of 10 typical imagers and 5 out of 10 aphantasic individuals showed positive FIN connectivity with the dorsolateral prefrontal cortex.

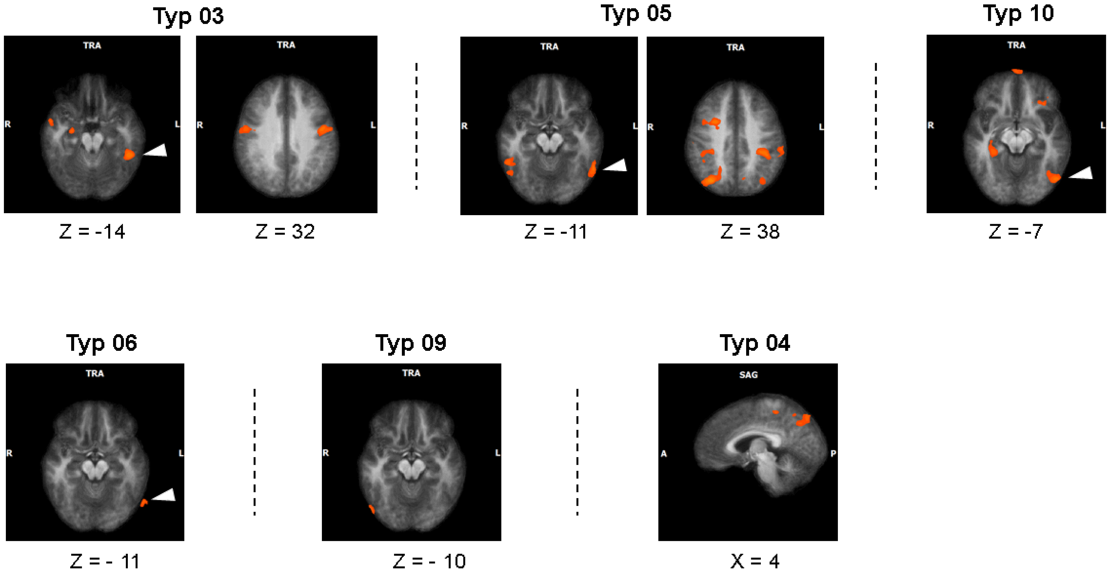


**Fig S11. Individual trial-by-trial parametric modulation of vividness in typical imager individuals (Typ).** Orange ROIs show significant clusters of which the activity during imagery is modulated by individual ratings of subjective vividness. Write arrows indicate the individual FIN. Seven out of 10 typical imagers rated vividness differently than the ceiling score in at least one trial per run. In 6 out of these 7 typical imagery individuals, we observed significant regions in which the activity was modulated by trial-by-trial vividness (data smoothed at 6mm, $p = 0.005$, number of voxel size > 150, arbitrary). The ROIs were projected to a group-averaging brain.

**Table S1. Coordinates for the imagery domain-general activation and domain-specific activation in typical imagers.** Cluster size thresholded at alpha=0.05, initial p=0.005, Monte-Carlo simulation n = 5000. Abs. = Abstract. P. = Perception.

| Condition | TAL X | TAL Y | TAL Z | Location | Size (mm3) | Average t | Average p | Peak t | Peak p |
|---|---|---|---|---|---|---|---|---|---|
| **all Imagery > Abs.** | -24 | -62 | 36 | L IPSp | 2584 | 4.40 | 0.0023 | 7.57 | 0.00003 |
| | -37 | -44 | 36 | L IPSa | 824 | 4.23 | 0.0026 | 5.36 | 0.00045 |
| | -41 | 31 | 17 | L IFG-Tri | 1683 | 4.32 | 0.0025 | 7.13 | 0.00006 |
| | -41 | -55 | -10 | L Fusi | 413 | 3.86 | 0.0040 | 5.89 | 0.00023 |
| **all imagery > all Perception** | 37 | -65 | -2 | R Visual area | 27368 | -6.71 | 0.0003 | -17.66 | 0.00000 |
| | 16 | -27 | 2 | R LGN | 1549 | -7.28 | 0.0002 | -22.13 | 0.00000 |
| | -16 | -26 | 0 | L LGN | 1006 | -6.03 | 0.0003 | -9.28 | 0.00001 |
| | -32 | -74 | -5 | L Visual area | 20530 | -7.06 | 0.0002 | -21.92 | 0.00000 |
| **Shape > Others** | 22 | -91 | 17 | R Occi. Sup | 724 | 3.92 | 0.0037 | 4.94 | 0.00080 |
| | -34 | -30 | -13 | L PHG | 764 | 4.75 | 0.0018 | 7.78 | 0.00003 |
| **Color > others** | -28 | -46 | -14 | L Fusi | 750 | 4.37 | 0.0024 | 7.13 | 0.00006 |
| **Word > others** | 55 | -48 | -6 | R pMTS | 2073 | 6.15 | 0.0003 | 10.32 | 0.00000 |
| | 47 | 10 | 32 | R IFG-Operc | 1328 | 5.46 | 0.0005 | 8.19 | 0.00002 |
| | 38 | -47 | 44 | R IPS | 7222 | 5.88 | 0.0004 | 11.16 | 0.00000 |
| | -47 | 4 | 29 | L IFG-Operc | 2834 | 5.70 | 0.0005 | 9.76 | 0.00000 |
| | -47 | -41 | 36 | L SMG | 2145 | 5.72 | 0.0004 | 9.21 | 0.00001 |
| | -52 | -58 | -4 | L Fusi | 2633 | 5.42 | 0.0005 | 7.58 | 0.00003 |
| **Face > others** | 58 | -5 | 2 | R mSTG | 612 | 4.85 | 0.0011 | 6.24 | 0.00015 |
| | 26 | -2 | -12 | R Amygdala | 3079 | 5.58 | 0.0006 | 10.93 | 0.00000 |
| | 4 | 61 | 16 | R vmPFC | 648 | 4.49 | 0.0017 | 5.83 | 0.00025 |
| | -20 | -4 | -11 | L Amygdala | 1964 | 5.51 | 0.0007 | 9.93 | 0.00000 |
| | -53 | -7 | -2 | L mSTG | 2391 | 4.85 | 0.0011 | 7.86 | 0.00003 |
| | -3 | -50 | 23 | L vPCC | 1915 | 4.88 | 0.0013 | 8.46 | 0.00001 |
| | -2 | 64 | 2 | L vmPFC | 629 | 4.75 | 0.0013 | 6.23 | 0.00015 |
| | -4 | 34 | -10 | L vmPFC | 649 | 4.31 | 0.0024 | 5.66 | 0.00031 |
| **Map > others** | 49 | -38 | 38 | R SMG | 761 | 6.08 | 0.0004 | 14.37 | 0.00000 |
| | 36 | -71 | 32 | R ANG | 1916 | 6.05 | 0.0003 | 10.37 | 0.00000 |
| | 28 | 3 | 51 | R PMd | 4056 | 5.97 | 0.0004 | 12.83 | 0.00000 |
| | 13 | -52 | 16 | R vPCC | 1931 | 5.77 | 0.0004 | 8.69 | 0.00001 |
| | 0 | -56 | 46 | L/R Precuneus | 6452 | 6.56 | 0.0003 | 20.40 | 0.00000 |
| | -14 | -54 | 13 | L vPCC | 1400 | 5.58 | 0.0005 | 8.55 | 0.00001 |
| | -26 | -2 | 50 | L PMd | 2503 | 5.62 | 0.0004 | 8.46 | 0.00001 |
| | -33 | -75 | 37 | L ANG | 826 | 5.17 | 0.0006 | 6.37 | 0.00013 |
| | -53 | -41 | 32 | L SMG | 2077 | 5.95 | 0.0004 | 13.54 | 0.00000 |
| | -26 | -35 | -8 | L PHG | 420 | 4.43 | 0.0022 | 6.91 | 0.00007 |

**Table S2. Domain-specific activation for perception in typical imagers.**

| Condition | TAL X | TAL Y | TAL Z | Location | Size (mm3) | Average t | Average p | Peak t | Peak p |
|---|---|---|---|---|---|---|---|---|---|
| **Shape > others** | 22 | -55 | -5 | R Fusi | 387 | 6.04 | 0.0004 | 9.88 | 0.00000 |
| | -27 | -49 | -11 | L FG | 443 | 5.62 | 0.0004 | 7.62 | 0.00003 |
| | -41 | -75 | -1 | L Occi. Inf | 736 | 5.42 | 0.0005 | 7.76 | 0.00003 |
| **Color > others** | -27 | -49 | -13 | L Fusi | 409 | 5.79 | 0.0004 | 8.46 | 0.00001 |
| | -32 | -23 | -17 | L PHG | 233 | 6.04 | 0.0004 | 9.88 | 0.00000 |
| **Word > others** | 56 | -49 | -8 | R Fusi | 598 | 5.68 | 0.0004 | 9.08 | 0.00001 |
| | -46 | -38 | 41 | L/R IPS | 14306 | 5.66 | 0.0005 | 10.05 | 0.00000 |
| | 32 | 44 | 23 | R AntFrontal | 672 | 5.84 | 0.0004 | 8.80 | 0.00001 |
| | 24 | -1 | 53 | R PMd | 2734 | 5.73 | 0.0004 | 10.13 | 0.00000 |
| | 19 | -90 | -2 | R Lingual | 731 | 5.78 | 0.0004 | 9.26 | 0.00001 |
| | -13 | -73 | 12 | L Calcarine | 639 | 5.55 | 0.0005 | 7.27 | 0.00005 |
| | -24 | -88 | -8 | L Lingual | 4212 | 6.24 | 0.0003 | 11.57 | 0.00000 |
| | -43 | -38 | 41 | L SMG | 4819 | 5.91 | 0.0004 | 11.82 | 0.00000 |
| | -26 | -4 | 54 | L PMd | 1252 | 5.85 | 0.0004 | 10.65 | 0.00000 |
| | -32 | 35 | 26 | L AntFrontal | 314 | 5.18 | 0.0006 | 5.83 | 0.00025 |
| | -46 | 1 | 39 | L PMv | 315 | 5.13 | 0.0007 | 6.06 | 0.00019 |
| | -49 | -59 | -7 | L Fusi | 1989 | 5.68 | 0.0004 | 9.05 | 0.00001 |
| **Face > others** | 41 | -80 | -8 | R Occi Inf | 278 | 5.54 | 0.0005 | 7.19 | 0.00005 |
| | 40 | -45 | -17 | R Fusi | 361 | 5.80 | 0.0004 | 7.84 | 0.00003 |
| | 19 | -7 | -11 | R Amygdala | 1048 | 6.51 | 0.0003 | 11.06 | 0.00000 |
| | -20 | -7 | -11 | L Amygdala | 778 | 6.74 | 0.0003 | 12.01 | 0.00000 |
| **Map > others** | 35 | -71 | 31 | R ANG | 1874 | 6.15 | 0.0003 | 10.85 | 0.00000 |
| | 17 | -53 | 18 | R vPCC | 1450 | 5.57 | 0.0005 | 9.29 | 0.00001 |
| | 21 | 1 | 50 | R PMd | 741 | 5.67 | 0.0004 | 8.48 | 0.00001 |
| | 0 | -64 | 47 | L/R Precuneus | 5681 | 6.80 | 0.0003 | 16.25 | 0.00000 |
| | -25 | -1 | 51 | L PMd | 1995 | 5.88 | 0.0004 | 10.54 | 0.00000 |

| | | | | | | | | | |
|---|---|---|---|---|---|---|---|---|---|
| | -16 | -59 | 20 | L vPCC | 272 | 5.41 | 0.0005 | 7.88 | 0.00003 |
| | -29 | -80 | 30 | L ANG | 1077 | 5.53 | 0.0005 | 8.14 | 0.00002 |
| | -61 | -50 | 8 | L pMTG | 269 | 5.70 | 0.0004 | 8.47 | 0.00001 |

**Table S3. Task-specific functional connectivity of the FIN in typical imagers during imagery.**

| Condition | TAL X | TAL Y | TAL Z | Location | Size (mm3) | Average t | Average p | Peak t | Peak p |
|---|---|---|---|---|---|---|---|---|---|
| **all Imagery > Abs.** | 24 | 34 | -6 | R OFC | 481 | 4.61 | 0.0021 | 7.56 | 0.00004 |
| | -31 | 35 | -5 | L OFC | 303 | 4.50 | 0.0021 | 7.10 | 0.00006 |
| **all Imagery > all Perception** | 37 | -71 | -1 | R Visual area | 21100 | -5.21 | 0.0015 | -17.3 | 0.00000 |
| | 46 | 10 | -14 | R ATL | 385 | 4.28 | 0.0026 | 6.03 | 0.00020 |
| | -35 | -77 | -1 | L Visual area | 15957 | -5.57 | 0.0012 | -18.1 | 0.00000 |
| | -31 | 33 | -8 | L OFC | 333 | 4.30 | 0.0024 | 5.62 | 0.00033 |
| | -53 | -8 | -4 | L mSTS | 963 | 4.68 | 0.0020 | 12.39 | 0.00000 |
| **Shape imagery > Abs.** | 52 | -28 | 35 | R SMG | 697 | 4.16 | 0.0029 | 6.28 | 0.00015 |
| | 56 | -58 | -2 | R Fusi | 975 | 4.62 | 0.0019 | 8.09 | 0.00002 |
| | 44 | 27 | 30 | R MFG | 365 | 4.32 | 0.0024 | 6.10 | 0.00018 |
| | 42 | -46 | 53 | R IPSa | 575 | 4.23 | 0.0027 | 6.07 | 0.00019 |
| | 37 | -69 | 31 | R ANG | 507 | 4.07 | 0.0031 | 5.18 | 0.00058 |
| | -23 | 53 | 6 | L Frontal pole | 540 | 4.18 | 0.0029 | 6.40 | 0.00013 |
| | -36 | -41 | 40 | L IPSa | 411 | 4.62 | 0.0021 | 8.51 | 0.00001 |
| | -49 | -58 | -5 | L Fusi | 917 | 4.89 | 0.0018 | 10.52 | 0.00000 |
| **Color imagery > Abs.** | 45 | -50 | -13 | R Fusi | 1018 | 4.37 | 0.0023 | 6.16 | 0.00017 |
| | 44 | 37 | 13 | R IFG-Tri | 1124 | 4.15 | 0.0029 | 6.54 | 0.00011 |
| | 43 | -50 | 35 | R SMG | 1245 | 4.63 | 0.0021 | 8.36 | 0.00002 |
| | 30 | -65 | 37 | R ANG | 931 | 4.43 | 0.0022 | 6.08 | 0.00018 |
| | 25 | 34 | -7 | R OFC | 794 | 4.96 | 0.0017 | 10.95 | 0.00000 |
| | -17 | 31 | 47 | L SFG | 1023 | 4.40 | 0.0023 | 6.06 | 0.00019 |
| | -45 | -62 | 37 | L ANG | 719 | 4.16 | 0.0028 | 5.65 | 0.00031 |
| | -59 | -30 | -10 | L MTG | 440 | 4.10 | 0.0029 | 5.11 | 0.00063 |
| **Word imagery > Abs.** | 52 | -55 | -11 | R Fusi | 419 | 4.48 | 0.0021 | 6.67 | 0.00009 |
| | 1 | -63 | 34 | L/R Precuneus | 610 | 4.05 | 0.0032 | 5.97 | 0.00021 |
| | -52 | -59 | -3 | L Fusi | 899 | 4.75 | 0.0020 | 11.91 | 0.00000 |
| **Face imagery > Abs.** | 44 | -50 | -10 | R Fusi | 3324 | 4.52 | 0.0022 | 8.26 | 0.00002 |
| | 44 | -72 | 32 | R ANG | 971 | 4.64 | 0.0020 | 7.83 | 0.00003 |
| | 43 | 32 | 16 | R IFS | 371 | 4.34 | 0.0024 | 6.43 | 0.00012 |
| | 25 | 2 | 50 | R PMd | 1047 | 4.39 | 0.0025 | 7.08 | 0.00006 |
| | 31 | 1 | -8 | R Amygdala | 453 | 4.50 | 0.0021 | 6.84 | 0.00008 |
| | 17 | 32 | -8 | R OFC | 1709 | 4.44 | 0.0023 | 8.67 | 0.00001 |
| | 16 | 21 | 49 | R SFG | 1925 | 4.51 | 0.0022 | 8.82 | 0.00001 |
| | 2 | 53 | 11 | R dmPFC | 1367 | 4.32 | 0.0026 | 7.22 | 0.00005 |
| | -7 | -48 | 22 | L vPCC | 1136 | 4.30 | 0.0026 | 7.67 | 0.00003 |
| | -17 | 25 | 44 | L PMd | 4920 | 4.57 | 0.0020 | 7.95 | 0.00002 |
| | -18 | 62 | 10 | L Frontal pole | 507 | 3.96 | 0.0035 | 4.66 | 0.00119 |
| | -22 | -4 | -10 | L Amygdala | 409 | 4.36 | 0.0024 | 5.90 | 0.00023 |

| | | | | | | | | | |
|---|---|---|---|---|---|---|---|---|---|
| | -34 | -39 | 37 | L IPS | 409 | 4.08 | 0.0030 | 5.05 | 0.00069 |
| | -37 | -16 | -16 | L PHG | 566 | 4.37 | 0.0022 | 6.37 | 0.00013 |
| | -50 | -18 | -13 | L MTG | 448 | 4.43 | 0.0022 | 6.74 | 0.00009 |
| | -53 | -58 | 34 | L ANG | 619 | 4.18 | 0.0027 | 5.37 | 0.00045 |
| **Map imagery > Abs.** | 26 | 12 | 52 | R PMd | 1385 | 4.21 | 0.0028 | 6.21 | 0.00016 |
| | 32 | -61 | 42 | R IPS | 626 | 3.96 | 0.0034 | 4.71 | 0.00110 |
| | -4 | -29 | 37 | L dPCC | 384 | 4.14 | 0.0029 | 5.78 | 0.00027 |
| | -23 | 22 | 43 | L SFS | 1892 | 4.20 | 0.0027 | 5.96 | 0.00021 |
| | -17 | 10 | 21 | L Caudate | 356 | 4.32 | 0.0025 | 7.02 | 0.00006 |
| | -46 | -61 | 34 | L ANG | 503 | 4.42 | 0.0023 | 6.47 | 0.00012 |
| | -35 | -17 | -16 | L PHG | 324 | 4.29 | 0.0025 | 6.18 | 0.00016 |

**Table S4. Task-specific functional connectivity of the FIN in typical imagers during perception.**

| Condition | TAL X | TAL Y | TAL Z | Location | Size (mm3) | Average t | Average p | Peak t | Peak p |
|---|---|---|---|---|---|---|---|---|---|
| **Shape Perception > Abs.** | 43 | -44 | -9 | R Fusi | 3701 | 4.77 | 0.0019 | 11.03 | 0.00000 |
| | 32 | -60 | 26 | R ANG | 982 | 4.29 | 0.0025 | 6.60 | 0.00010 |
| | 26 | 15 | 46 | R FEF | 342 | 4.26 | 0.0027 | 7.78 | 0.00003 |
| | -24 | 11 | 44 | L FEF | 364 | 4.47 | 0.0025 | 10.29 | 0.00000 |
| | -41 | -48 | -7 | L Fusi | 528 | 4.35 | 0.0025 | 6.39 | 0.00013 |
| **Color perception > Abs.** | 33 | -62 | -8 | R Fusi | 6307 | 4.74 | 0.0029 | 10.71 | 0.00000 |
| | 45 | -42 | 29 | R ANG | 464 | 4.48 | 0.0030 | 6.70 | 0.00009 |
| | 22 | 15 | 48 | R PMd | 829 | 4.41 | 0.0024 | 8.25 | 0.00002 |
| | 26 | -53 | 33 | R ANG | 725 | 4.13 | 0.0026 | 5.86 | 0.00024 |
| | 26 | 45 | -2 | R AntFrontal | 298 | 4.15 | 0.0030 | 5.39 | 0.00044 |
| | 6 | 27 | 33 | R FEF | 314 | 4.53 | 0.0024 | 6.19 | 0.00016 |
| | -29 | 48 | 1 | L AntFrontal | 490 | 4.47 | 0.0028 | 8.95 | 0.00001 |
| | -20 | -73 | -9 | L Lingual | 625 | 4.69 | 0.0023 | 7.24 | 0.00005 |
| | -33 | -58 | -8 | L Fusi | 2767 | 4.56 | 0.0012 | 13.39 | 0.00000 |
| **Word perception > Abs.** | 33 | -50 | -6 | R Fusi | 20908 | 4.64 | 0.0008 | 15.03 | 0.00000 |
| | 30 | 9 | 46 | R PMd | 923 | 4.16 | 0.0028 | 6.24 | 0.00015 |
| | -16 | -59 | 37 | R IPS | 423 | 4.47 | 0.0027 | 6.46 | 0.00012 |
| | -39 | -47 | -6 | L Fusi | 7160 | 4.62 | 0.0026 | 8.49 | 0.00001 |
| | -31 | -32 | 26 | L SMG | 603 | 4.46 | 0.0027 | 6.85 | 0.00008 |
| | -42 | -48 | 25 | L ANG | 488 | 4.37 | 0.0026 | 5.73 | 0.00029 |

| | | | | | | | | | |
|---|---|---|---|---|---|---|---|---|---|
| **Face perception > Abs.** | 33 | -49 | -7 | R Fusi | 12405 | 4.68 | 0.0008 | 16.11 | 0.00000 |
| | 37 | 25 | 12 | R IFS | 1431 | 4.53 | 0.0024 | 7.97 | 0.00002 |
| | 39 | -59 | 25 | R ANG | 427 | 4.69 | 0.0026 | 8.10 | 0.00002 |
| | 26 | 9 | 40 | R FEF | 1416 | 4.69 | 0.0027 | 8.49 | 0.00001 |
| | 27 | -19 | -16 | R PHG | 361 | 4.10 | 0.0029 | 5.77 | 0.00027 |
| | 26 | -36 | 32 | R SMG | 312 | 4.54 | 0.0029 | 6.76 | 0.00008 |
| | 20 | -55 | 31 | R dPCC | 428 | 4.20 | 0.0023 | 5.63 | 0.00032 |
| | 18 | 27 | -5 | R OFC | 395 | 4.36 | 0.0030 | 5.80 | 0.00026 |
| | -45 | -49 | -3 | L Fusi | 2651 | 4.32 | 0.0030 | 7.55 | 0.00004 |
| | -31 | -36 | -7 | L PHG | 564 | 4.34 | 0.0027 | 7.27 | 0.00005 |
| **Map perception > Abs.** | 35 | -65 | -10 | R Fusi | 9552 | 4.84 | 0.0015 | 12.47 | 0.00000 |
| | 40 | -58 | 24 | R ANG | 5627 | 4.63 | 0.0023 | 8.55 | 0.00001 |
| | 40 | 42 | 9 | R MFG | 305 | 4.19 | 0.0029 | 5.29 | 0.00050 |
| | 32 | 10 | 44 | R FEF | 1088 | 4.53 | 0.0027 | 6.72 | 0.00009 |
| | 12 | 36 | 35 | R FEF | 502 | 4.15 | 0.0028 | 6.22 | 0.00016 |
| | 7 | -51 | 24 | R vPCC | 307 | 4.80 | 0.0027 | 7.76 | 0.00003 |
| | -1 | -31 | 30 | L vPCC | 1531 | 4.76 | 0.0016 | 12.18 | 0.00000 |
| | -5 | 29 | 5 | L vACC | 807 | 4.36 | 0.0030 | 7.62 | 0.00003 |
| | -16 | 32 | 36 | L FEF | 310 | 4.10 | 0.0028 | 6.36 | 0.00013 |
| | -32 | 14 | 31 | L FEF | 734 | 4.29 | 0.0025 | 6.33 | 0.00014 |
| | -45 | -47 | -4 | L Fusi | 267 | 4.47 | 0.0027 | 9.69 | 0.00001 |
| | -31 | -35 | -7 | L PHG | 2302 | 4.15 | 0.0025 | 5.79 | 0.00026 |
| | -43 | -47 | 28 | L ANG | 1784 | 4.68 | 0.0025 | 6.68 | 0.00009 |
| | -47 | -23 | 32 | L SMG | 445 | 4.35 | 0.0023 | 6.32 | 0.00014 |

**Table S5. Imagery domain-general activation and domain-specific activation in aphantasics.**

| Condition | TAL X | TAL Y | TAL Z | Location | Size (mm3) | Average t | Average p | Peak t | Peak p |
|---|---|---|---|---|---|---|---|---|---|
| **all Imagery > Abs. (p<0.001)** | 55 | -22 | 36 | R SMG | 360 | 5.20 | 0.0006 | 6.32 | 0.00014 |
| | 50 | 8 | 35 | R PMd | 2158 | 5.64 | 0.0004 | 11.06 | 0.00000 |
| | 39 | -40 | 43 | R IPS | 1148 | 5.25 | 0.0006 | 6.86 | 0.00007 |
| | 30 | 0 | 52 | R PMv | 2928 | 5.68 | 0.0005 | 10.79 | 0.00000 |
| | 30 | -62 | 42 | R ANG | 479 | 5.14 | 0.0007 | 6.11 | 0.00018 |
| | -5 | -5 | 49 | L Supp. motor | 288 | 5.12 | 0.0007 | 5.98 | 0.00021 |
| | -32 | -53 | 41 | L ANG | 6403 | 5.81 | 0.0004 | 11.25 | 0.00000 |
| | -27 | -8 | 50 | L PMd | 1288 | 5.32 | 0.0006 | 7.43 | 0.00004 |

| | | | | | | | | | |
|---|---|---|---|---|---|---|---|---|---|
| | -37 | 37 | 13 | L IFG-Tri | 381 | 5.10 | 0.0007 | 5.82 | 0.00025 |
| | -47 | 3 | 34 | L PMv | 1266 | 5.41 | 0.0005 | 7.50 | 0.00004 |
| | -43 | 26 | 27 | L IFG-Tri | 916 | 5.70 | 0.0004 | 8.61 | 0.00001 |
| | -45 | -54 | -10 | L Fusi | 514 | 5.90 | 0.0004 | 9.22 | 0.00001 |
| | -48 | 42 | -5 | L IFG-Orb | 422 | 5.71 | 0.0004 | 8.37 | 0.00002 |
| | -47 | 44 | 15 | L IFG-Tri | 494 | 5.61 | 0.0005 | 7.97 | 0.00002 |
| | -47 | -31 | 21 | L SMG | 274 | 5.76 | 0.0004 | 8.04 | 0.00002 |
| | -64 | -28 | 27 | L IPS | 555 | 5.30 | 0.0006 | 6.80 | 0.00008 |
| **all Imagery vs P.** | 35 | -68 | -7 | R Visual area | 22230 | -5.21 | 0.0014 | -10.95 | 0.00000 |
| | 17 | -27 | 2 | R LGN | 1012 | -4.70 | 0.0018 | -7.26 | 0.00005 |
| | 2 | -62 | 0 | L/R EVA | 5454 | 4.77 | 0.0018 | 10.80 | 0.00000 |
| | -1 | -85 | 22 | L/R EVA | 5401 | 4.34 | 0.0024 | 6.87 | 0.00007 |
| | 3 | 1 | 55 | L/R Supp.motor | 732 | 4.32 | 0.0024 | 5.68 | 0.00030 |
| | -19 | -27 | 0 | L LGN | 703 | -4.75 | 0.0018 | -7.36 | 0.00004 |
| | -32 | -74 | -7 | L Visual area | 17515 | -5.28 | 0.0014 | -14.62 | 0.00000 |
| | -36 | -26 | 50 | L Prim. motor | 1850 | 4.44 | 0.0024 | 9.55 | 0.00001 |
| | -41 | -68 | 31 | L ANG | 1143 | -4.09 | 0.0030 | -4.83 | 0.00094 |
| | -53 | 6 | 13 | L IFG-Operc | 848 | 4.58 | 0.0021 | 8.03 | 0.00002 |
| **Color > Others** | 22 | 32 | -7 | R OFC | 401 | 4.67 | 0.0017 | 6.16 | 0.00017 |
| | -25 | 31 | -5 | L OFC | 514 | 4.31 | 0.0025 | 6.69 | 0.00009 |
| | -41 | 31 | 19 | L IFS | 1282 | 4.38 | 0.0023 | 6.44 | 0.00012 |
| **Word > Others** | 35 | -46 | 42 | R IPS | 1506 | 4.15 | 0.0029 | 5.61 | 0.00033 |
| | 37 | 39 | 28 | R MFG | 1922 | 4.85 | 0.0018 | 9.29 | 0.00001 |
| | 23 | -4 | 52 | R PMd | 1486 | 4.43 | 0.0022 | 6.74 | 0.00009 |
| | -24 | -63 | 46 | L IPS | 3086 | 4.35 | 0.0024 | 7.89 | 0.00003 |
| **Face > Others** | 49 | 5 | -4 | R STG | 442 | 5.92 | 0.0004 | 8.21 | 0.00002 |
| | 23 | -8 | -11 | R Amygdala | 1544 | 6.90 | 0.0003 | 15.24 | 0.00000 |
| | -6 | 52 | 29 | L dmPFC | 2427 | 5.84 | 0.0004 | 14.66 | 0.00000 |

| | | | | | | | | | |
|---|---|---|---|---|---|---|---|---|---|
| | -1 | 48 | -4 | L vmPFC | 2250 | 5.60 | 0.0005 | 9.34 | 0.00001 |
| | -2 | -57 | 27 | L vPCC | 637 | 5.20 | 0.0006 | 6.83 | 0.00008 |
| | -24 | -5 | -11 | L Amygdala | 2229 | 6.17 | 0.0003 | 12.11 | 0.00000 |
| | -52 | -10 | -2 | L STG | 3249 | 5.74 | 0.0005 | 10.95 | 0.00000 |
| **Map > Others** | 34 | -79 | 29 | R ANG | 1409 | 4.99 | 0.0012 | 7.80 | 0.00003 |
| | 27 | 2 | 50 | R PMd | 2768 | 4.57 | 0.0016 | 6.76 | 0.00008 |
| | 19 | -32 | -9 | R PHG | 2067 | 5.05 | 0.0011 | 9.62 | 0.00001 |
| | 1 | -62 | 46 | L/R Precuneus | 10083 | 5.18 | 0.0010 | 9.48 | 0.00001 |
| | -2 | -49 | 8 | L/R vPCC | 9598 | 5.25 | 0.0011 | 14.12 | 0.00000 |
| | -26 | 0 | 50 | L PMd | 2431 | 4.85 | 0.0013 | 8.74 | 0.00001 |
| | -29 | -82 | 30 | L ANG | 944 | 4.50 | 0.0017 | 6.68 | 0.00009 |

Abs = Abstract words; P = Perception

**Table S6. Perceptual domain-specific activation in aphantasics.**

| Condition | TAL X | TAL Y | TAL Z | Location | Size (mm3) | Average t | Average p | Peak t | Peak p |
|---|---|---|---|---|---|---|---|---|---|
| **Shape > others** | 45 | -69 | -2 | R Fusi | 298 | 5.37 | 0.0006 | 6.37 | 0.00013 |
| | -37 | -74 | -7 | L Fusi | 403 | 5.27 | 0.0008 | 6.26 | 0.00015 |
| **Color > others** | 55 | -22 | 10 | R STG | 512 | 5.45 | 0.0005 | 7.94 | 0.00002 |
| | 27 | -61 | -13 | R Fusi | 587 | 5.25 | 0.0008 | 8.46 | 0.00001 |
| | -4 | 49 | 15 | L vmPFC | 1091 | 5.22 | 0.0006 | 6.94 | 0.00007 |
| | -2 | -19 | 44 | L/R dPCC | 817 | 5.57 | 0.0005 | 8.18 | 0.00002 |
| | -28 | -34 | 53 | L Precentral | 361 | 5.26 | 0.0006 | 6.61 | 0.00010 |
| | -41 | 19 | -8 | L TempPole | 269 | 5.23 | 0.0006 | 6.61 | 0.00010 |
| | -47 | 35 | 16 | L dlPFC | 420 | 6.04 | 0.0004 | 10.75 | 0.00000 |
| | -62 | -26 | 13 | L STG | 699 | 6.10 | 0.0003 | 10.05 | 0.00000 |
| | -1 | -71 | 13 | L/R Calcarine | 1400 | 5.33 | 0.0006 | 7.66 | 0.00003 |

| | | | | | | | | | |
|---|---|---|---|---|---|---|---|---|---|
| | -27 | -46 | -14 | L Fusi | 293 | 5.32 | 0.0006 | 7.34 | 0.00004 |
| **Word > others** | 41 | -34 | 43 | R IPS | 247 | 5.36 | 0.0006 | 7.30 | 0.00005 |
| | -22 | -89 | -14 | L Lingual | 475 | 5.58 | 0.0005 | 8.96 | 0.00001 |
| | -20 | -66 | 44 | L SPL | 280 | 5.90 | 0.0004 | 8.99 | 0.00001 |
| | -37 | -34 | 35 | L IPS | 335 | 5.14 | 0.0006 | 5.99 | 0.00020 |
| | -40 | -63 | -8 | L Fusi | 230 | 5.04 | 0.0007 | 4.90 | 0.00085 |
| **Face > others** | 39 | -79 | -11 | R Occi Inf | 466 | 5.32 | 0.0006 | 7.14 | 0.00005 |
| | 20 | -5 | -8 | R Amygdala | 1351 | 5.20 | 0.0009 | 6.62 | 0.00010 |
| | 38 | -47 | -17 | R Fusi | 380 | 5.22 | 0.0008 | 5.41 | 0.00043 |
| | -1 | 56 | -2 | L vmPFC | 306 | 5.20 | 0.0008 | 7.67 | 0.00003 |
| | -3 | -47 | 26 | L vPCC | 624 | 5.32 | 0.0008 | 7.79 | 0.00003 |
| | -17 | -3 | -11 | L Amygdala | 311 | 5.00 | 0.0009 | 6.59 | 0.00010 |
| **Map > others** | 35 | -74 | 31 | R ANG | 3711 | 5.62 | 0.0005 | 12.01 | 0.00000 |
| | 29 | 1 | 54 | R PMd | 787 | 5.29 | 0.0006 | 6.53 | 0.00011 |
| | 1 | -67 | 47 | L/R Precuneus | 4839 | 5.48 | 0.0005 | 8.52 | 0.00001 |
| | 12 | -55 | 17 | R vPCC | 583 | 5.62 | 0.0004 | 7.31 | 0.00005 |
| | -30 | -2 | 55 | L PMd | 313 | 5.06 | 0.0007 | 5.98 | 0.00021 |
| | -36 | -69 | 38 | L ANG | 664 | 5.00 | 0.0008 | 6.90 | 0.00007 |
| | -22 | -35 | -8 | L PHG | 362 | 4.97 | 0.0009 | 7.65 | 0.00003 |

**Table S7. Task-specific functional connectivity of the FIN in aphantasics during imagery and perceptual domains.**

| Condition | TAL X | TAL Y | TAL Z | Location | Size (mm3) | Average t | Average p | Peak t | Peak p |
|---|---|---|---|---|---|---|---|---|---|
| **Color imagery > Abs.** | 44 | -56 | -13 | R Fusi | 386 | 4.31 | 0.0024 | 5.95 | 0.00022 |
| | -28 | -47 | -14 | L Fusi | 608 | 4.34 | 0.0025 | 7.28 | 0.00005 |
| **Map imagery > Abs.** | 11 | -52 | 14 | R vPCC | 824 | 4.44 | 0.0022 | 8.32 | 0.00002 |

| | | | | | | | | | |
|---|---|---|---|---|---|---|---|---|---|
| **Shape perception > Abs.** | 35 | -47 | -10 | R Fusi | 1737 | 4.37 | 0.0024 | 8.05 | 0.00002 |
| | 30 | -68 | 1 | R Lingual | 312 | 4.02 | 0.0032 | 4.73 | 0.00107 |
| | -29 | -54 | -3 | L Fusi | 535 | 4.19 | 0.0027 | 5.82 | 0.00025 |
| | -24 | -65 | -15 | L Fusi | 692 | 4.53 | 0.0027 | 6.32 | 0.00014 |
| | -29 | -66 | -1 | L Fusi | 403 | 4.16 | 0.0028 | 5.36 | 0.00046 |
| **Color perception > Abs.** | 34 | -46 | -13 | R Fusi | 908 | 4.77 | 0.0030 | 8.81 | 0.00001 |
| | 34 | -57 | -1 | R Fusi | 311 | 4.65 | 0.0026 | 8.24 | 0.00002 |
| | -31 | -64 | -4 | L Fusi | 737 | 4.33 | 0.0028 | 6.20 | 0.00016 |
| **Word perception > Abs.** | 36 | -38 | -15 | R Fusi | 1199 | 4.31 | 0.0030 | 5.92 | 0.00022 |
| | 28 | 38 | 1 | R AntFrontal | 1000 | 4.38 | 0.0028 | 6.02 | 0.00020 |
| | 30 | -67 | 14 | R Calcarine | 599 | 4.31 | 0.0028 | 7.24 | 0.00005 |
| | 19 | 51 | 8 | R AntFrontal | 863 | 5.29 | 0.0023 | 11.68 | 0.00000 |
| **Face perception > Abs.** | 28 | -56 | -9 | R Fusi | 1170 | 4.93 | 0.0029 | 12.48 | 0.00000 |
| | 29 | -67 | 6 | R Fusi | 1012 | 4.57 | 0.0027 | 7.06 | 0.00006 |
| | -32 | -65 | -5 | L Lingual | 2320 | 4.70 | 0.0022 | 12.15 | 0.00000 |

No significant area was identified in the imagery of shape, letter and face tasks, or in the map perception task.

**Table S8. Group differences of brain activations in imagery and in perception tasks.**

| Condition | TAL X | TAL Y | TAL Z | Location | Size (mm3) | Average t | Average p | Peak t | Peak p |
|---|---|---|---|---|---|---|---|---|---|
| **all Imagery AP > TI** | 56 | -53 | 4 | R pMTG | 1593 | 14.87 | 0.0018 | 26.23 | 0.00007 |
| | 58 | -26 | 35 | R SMG | 429 | 12.91 | 0.0026 | 19.96 | 0.00030 |
| | 47 | 11 | 8 | R IFG-Operc | 389 | 12.32 | 0.0028 | 16.87 | 0.00066 |
| | 46 | 46 | 0 | R IFG-Orb | 1636 | 14.94 | 0.0018 | 27.43 | 0.00006 |
| | 41 | -43 | 41 | R IPS | 266 | 12.76 | 0.0026 | 17.57 | 0.00055 |

| | | | | | | | | | |
|---|---|---|---|---|---|---|---|---|---|
| | -26 | -86 | -7 | L Lingual | 223 | 14.11 | 0.0021 | 22.42 | 0.00017 |
| | -31 | -79 | 2 | L Occi. Mid | 304 | 14.11 | 0.0021 | 24.49 | 0.00010 |
| **all Perception AP > TI** | 62 | -24 | 37 | R SMG | 389 | 12.84 | 0.0025 | 19.29 | 0.00035 |
| | 50 | 40 | -4 | R IFG-Orb | 323 | 12.71 | 0.0027 | 20.96 | 0.00023 |
| **Face perception TI > AP** | 61 | -24 | 38 | R SMG | 387 | -3.22 | 0.0023 | -3.71 | 0.00039 |
| | 49 | 37 | -6 | R IFG-Orb | 255 | -3.16 | 0.0026 | -3.69 | 0.00042 |
| | 44 | -65 | 17 | R pMTG | 400 | 3.37 | 0.0018 | 4.18 | 0.00008 |
| | 11 | -92 | -12 | R Lingual | 892 | 3.28 | 0.0020 | 4.07 | 0.00012 |
| | -36 | -59 | -11 | L Fusi | 863 | 3.61 | 0.0014 | 5.42 | 0.00000 |

**Table S9. Task-specific functional connectivity of the IFG-SMG network in aphantasics.**

| Condition | TAL X | TAL Y | TAL Z | Location | Size (mm3) | Average t | Average p | Peak t | Peak p |
|---|---|---|---|---|---|---|---|---|---|
| **all Imagery > Abs.** | 62 | -9 | 7 | R mSTG | 176 | 3.95 | 0.0034 | 4.37 | 0.00180 |
| | -42 | -35 | 16 | L pSTG | 1829 | 4.38 | 0.0024 | 7.97 | 0.00002 |
| | -62 | -40 | 5 | L MTG | 1203 | 4.37 | 0.0025 | 10.29 | 0.00000 |
| **all Perception > Abs.** | -16 | -97 | 15 | R Occi Sup | 1289 | 4.77 | 0.0018 | 8.67 | 0.00001 |
| | -40 | 39 | -5 | L IFG-Orb | 540 | 4.50 | 0.0020 | 6.01 | 0.00020 |
| | -42 | 3 | 43 | L PMd | 365 | 3.94 | 0.0035 | 4.33 | 0.00190 |
| | -57 | -56 | -2 | L Fusi | 1034 | 4.27 | 0.0026 | 7.21 | 0.00005 |
| | -59 | -27 | -7 | L MTG | 875 | 4.09 | 0.0030 | 5.36 | 0.00045 |

**Table S10. Group differences of task-specific functional connectivity of the FIN during imagery.**

| Condition | TAL X | TAL Y | TAL Z | Location | Size (mm3) | Average t | Average p | Peak t | Peak p |
|---|---|---|---|---|---|---|---|---|---|

| | | | | | | | | | |
|---|---|---|---|---|---|---|---|---|---|
| **all Imagery > Abs.** | -23 | 54 | 8 | L AntFrontal | 1109 | 13.70 | 0.0022 | 21.98 | 0.00018 |
| | -28 | 36 | -2 | L OFC | 403 | 13.25 | 0.0025 | 22.75 | 0.00015 |
| | -46 | -26 | -8 | L MTG | 485 | 14.41 | 0.0022 | 39.36 | 0.00001 |
| **all Imagery > all Perception** | -18 | 55 | 9 | L AntFrontal | 377 | 11.65 | 0.0033 | 14.63 | 0.00124 |
| | -31 | 37 | -6 | L OFC | 628 | 13.09 | 0.0025 | 22.41 | 0.00017 |
| | -59 | -8 | -2 | L STG | 228 | 12.18 | 0.0030 | 16.95 | 0.00065 |
| **Shape imagery > Abs.** | 25 | -71 | -16 | R Lingual | 861 | 14.88 | 0.0015 | 24.40 | 0.00011 |
| | -19 | 13 | 48 | L PMd | 422 | 16.48 | 0.0012 | 29.85 | 0.00003 |
| | -24 | 53 | 9 | L AntFrontal | 841 | 15.54 | 0.0013 | 28.91 | 0.00004 |
| | -31 | 41 | 28 | L dlPFC | 400 | 14.00 | 0.0017 | 21.52 | 0.00020 |
| | -28 | 13 | -2 | L Insula | 262 | 16.16 | 0.0013 | 33.14 | 0.00002 |
| | -40 | -23 | -7 | L MTG | 1101 | 16.55 | 0.0012 | 34.86 | 0.00001 |
| | -37 | 49 | 10 | L AntFrontal | 254 | 14.06 | 0.0018 | 23.74 | 0.00012 |
| **Color imagery > Abs.** | -28 | 4 | 48 | L PMd | 182 | 14.17 | 0.0020 | 20.81 | 0.00024 |
| | -38 | -19 | -8 | L MTG | 919 | 14.52 | 0.0020 | 29.46 | 0.00004 |
| **Word imagery > Abs.** | -2 | -59 | 35 | L/R Precuneus | 2390 | 17.19 | 0.0011 | 31.98 | 0.00002 |
| | -20 | 13 | 47 | L PMd | 255 | 14.69 | 0.0016 | 23.28 | 0.00014 |
| | -26 | 51 | 8 | L AntFrontal | 891 | 16.92 | 0.0013 | 47.23 | 0.00000 |
| | -47 | -62 | 32 | L ANG | 433 | 13.07 | 0.0021 | 16.70 | 0.00069 |
| | -44 | -25 | -9 | L MTG | 641 | 16.12 | 0.0012 | 28.77 | 0.00004 |
| **Face imagery > Abs.** | 48 | -51 | -9 | R ITG | 449 | 12.31 | 0.0030 | 27.86 | 0.00005 |
| | -5 | 31 | 34 | L FEF | 314 | 12.97 | 0.0024 | 18.20 | 0.00047 |
| | -22 | 11 | 46 | L PMd | 817 | 14.80 | 0.0020 | 35.28 | 0.00001 |
| | -37 | 49 | 7 | L AntFrontal | 713 | 13.84 | 0.0022 | 26.86 | 0.00006 |
| | -40 | -20 | -10 | L MTG | 678 | 14.17 | 0.0021 | 26.47 | 0.00007 |
| **Map imagery > Abs.** | -20 | 0 | 37 | L PMd | 189 | 12.24 | 0.0029 | 16.71 | 0.00069 |
| | -40 | -20 | -10 | L MTG | 470 | 13.20 | 0.0024 | 22.92 | 0.00015 |

**Table S11. Group differences of task-specific functional connectivity of the FIN during perception.**

| Condition | TAL X | TAL Y | TAL Z | Location | Size (mm3) | Average t | Average p | Peak t | Peak p |
|---|---|---|---|---|---|---|---|---|---|
| **all Perception > Abs.** | -9 | -100 | 8 | R FG | 204 | 11.90 | 0.0031 | 15.76 | 0.00078 |
| **Shape Perception > Abs.** | -20 | 13 | 47 | L PMd | 256 | 14.49 | 0.0021 | 28.14 | 0.00005 |
| **Color Perception > Abs.** | 46 | -14 | 42 | R Prim. motor | 321 | 14.97 | 0.0017 | 23.89 | 0.00012 |
| | -10 | 31 | 31 | L FEF | 178 | 13.14 | 0.0025 | 20.47 | 0.00026 |
| | -40 | -19 | -11 | L MTG | 575 | 13.98 | 0.0021 | 23.97 | 0.00012 |

| | | | | | | | | | |
|---|---|---|---|---|---|---|---|---|---|
| **Word Perception > Abs.** | 42 | -49 | -12 | R OTS | 257 | 11.22 | 0.0048 | 18.61 | 0.00042 |
| | 14 | -51 | 49 | R SPL | 175 | 11.25 | 0.0045 | 16.29 | 0.00078 |
| | -20 | 11 | 47 | L PMd | 243 | 10.81 | 0.0050 | 16.35 | 0.00076 |
| | -41 | -23 | 1 | L STS | 230 | 11.75 | 0.0040 | 18.82 | 0.00040 |
| **Face Perception > Abs.** | 50 | -65 | 14 | R LOTC | 362 | 12.18 | 0.0029 | 15.98 | 0.00084 |
| | 41 | -47 | 13 | R OTS | 523 | 12.16 | 0.0030 | 19.85 | 0.00031 |
| | 40 | -72 | -13 | R OTS | 284 | 12.22 | 0.0029 | 16.27 | 0.00078 |
| | 28 | -3 | -5 | R Amygdala | 407 | 12.88 | 0.0027 | 22.75 | 0.00015 |
| | -13 | -7 | -10 | L Amygdala | 354 | 14.00 | 0.0024 | 33.97 | 0.00002 |
| | -21 | 3 | 40 | L PMd | 1481 | 14.64 | 0.0020 | 37.41 | 0.00001 |
| | -38 | -19 | -10 | L MTG | 1000 | 15.52 | 0.0018 | 35.56 | 0.00001 |
| **Map Perception > Abs.** | -25 | 10 | 45 | L PMd | 462 | 12.60 | 0.0028 | 20.15 | 0.00028 |
| | -48 | -26 | -10 | L MTG | 1782 | 16.42 | 0.0017 | 54.12 | 0.00000 |
